# Control of G protein-coupled receptor function via membrane-interacting intrinsically disordered C-terminal domains

**DOI:** 10.1101/2023.08.16.553551

**Authors:** Chiara Mancinelli, Dagan C. Marx, Alberto J. Gonzalez-Hernandez, Kevin Huynh, Lucia Mancinelli, Anisul Arefin, George Khelashvilli, Joshua Levitz, David Eliezer

## Abstract

G protein-coupled receptors (GPCRs) control intracellular signaling cascades via agonist-dependent coupling to intracellular transducers including heterotrimeric G proteins, GPCR kinases (GRKs), and arrestins. In addition to their critical interactions with the transmembrane core of active GPCRs, all three classes of transducers have also been reported to interact with receptor C-terminal domains (CTDs). An underexplored aspect of GPCR CTDs is their possible role as lipid sensors given their proximity to the membrane. CTD-membrane interactions have the potential to control the accessibility of key regulatory CTD residues to downstream effectors and transducers. Here we report that the CTDs of two closely related family C GPCRs, metabotropic glutamate receptor 2 (mGluR2) and mGluR3, bind to membranes and that this interaction can regulate receptor function. We first characterize CTD structure with NMR spectroscopy, revealing lipid composition-dependent modes of membrane binding. Using molecular dynamics simulations and structure-guided mutagenesis, we then identify key conserved residues and cancer-associated mutations that modulate CTD-membrane binding. Finally, we provide evidence that mGluR3 transducer coupling is controlled by CTD-membrane interactions in live cells, which may be subject to regulation by CTD phosphorylation and changes in membrane composition. This work reveals a novel mechanism of GPCR modulation, suggesting that CTD-membrane binding may be a general regulatory mode throughout the broad GPCR superfamily.

**Significance Statement:** G protein-coupled receptors (GPCRs) allow cells to sense and respond to their environment and constitute the largest class of targets for approved therapeutic drugs. Temporally precise GPCR signaling is achieved by coupling the binding of extracellular ligands to the binding of intracellular signal transducers (e.g. heterotrimeric G proteins) and regulators (e.g. β-arrestins). The C-terminal domains (CTDs) of GPCRs are targets of various post-translational modifications and play a critical role in transducer and regulator recruitment. Here we report novel interactions of the CTDs of two GPCRs of the metabotropic glutamate receptor family with cellular membranes. These interactions serve to regulate CTD accessibility and thus, mGluR coupling to transducers and regulators. We propose that dynamic CTD-membrane interaction constitutes a general mechanism for regulating GPCR function.

## Introduction

G protein-coupled receptors (GPCRs) respond to extracellular stimuli to drive intracellular signal transduction pathways that control a wide variety of biological functions. Consistent with their widespread physiological roles, GPCRs also serve as a major class of targets for disease intervention^1,2^. All GPCRs share a conserved architecture including an N-terminal extracellular domain (ECD) of variable size, a seven-helix transmembrane domain (TMD) and an intracellular C-terminal domain (CTD). Signaling is initiated by binding of extracellular ligands to the receptor ECD and/or TMD, inducing conformational changes that control coupling of receptor TMD and CTD to intracellular transducers, including heterotrimeric G proteins, GPCR kinases (GRKs) and β-arrestins (β-arrs).

GPCR CTDs typically feature low sequence complexity, are absent from most structures determined by X-ray crystallography or cryo-EM, and usually lack secondary structure in sequence-based predictions, including AlphaFold, suggesting that they are, in general, highly dynamic. Indeed, several recent studies of isolated or unbound GPCR CTDs have shown that they are intrisically disordered^3–6^. Well-defined GPCR CTD conformations have been captured at high resolution in only a handful of complexes that feature CTD-G protein^7–10^ or CTD-arrestin^11–17^ interactions and are typically limited to small segments. In addition to their roles in direct interaction with transducers, the proximity of GPCR CTDs to the membrane may also promote direct interactions with phospholipids. Indeed, membrane binding has been observed for the isolated CTD of the cannabinoid receptor 1^18^, and many GPCR CTDs are palmitoylated^19^, but the functional implications of CTD-membrane interactions are unclear. CTD-membrane interactions could influence receptor interactions and conformation and thereby modulate receptor ligand binding, activation, and trafficking. Furthermore, such interactions could be sensitive to lipid composition, providing one avenue by which lipids could dynamically regulate receptor function. Along these lines, recent work has demonstrated that reconstituted GPCR activity can be tuned by interactions with specific lipids^20–22^, and that β-arr can also directly interact with the lipid bilayer in its receptor-bound state^16,23–25^.

Metabotropic glutamate receptors (mGluRs) are dimeric, family C GPCRs that are characterized structurally by their large ECDs which contain a ligand binding domain (LBD) that senses the neurotransmitter glutamate and a cysteine-rich domain that connects the LBD to the TMD (**Fig. 1A**)^26^. Despite this unique ECD arrangement, upon glutamate binding by the LBD, mGluRs couple to G proteins via their TMD in a manner generally analogous to, yet distinct in detail from, that of other GPCR families^9,27–29^. As with other GPCRs, the CTDs of mGluRs are known to be major determinants of their interactions with transducers and regulatory factors^30–33^. Notably, we recently found that modest differences in CTD composition control the ability of the highly homologous group II mGluRs, mGluR2 and mGluR3, to recruit β-arrs^34^. mGluR3 is efficiently phosphorylated by GRKs and recruits β-arrs, which initiate clathrin-mediated receptor endocytosis, while mGluR2 largely eludes β-arrs driven internalization. This difference is encoded in a short ∼20 residue serine/threonine (S/T) rich region of the CTD that begins ∼15 residues after the end of TMD helix 7^34^.

**Figure 1:**
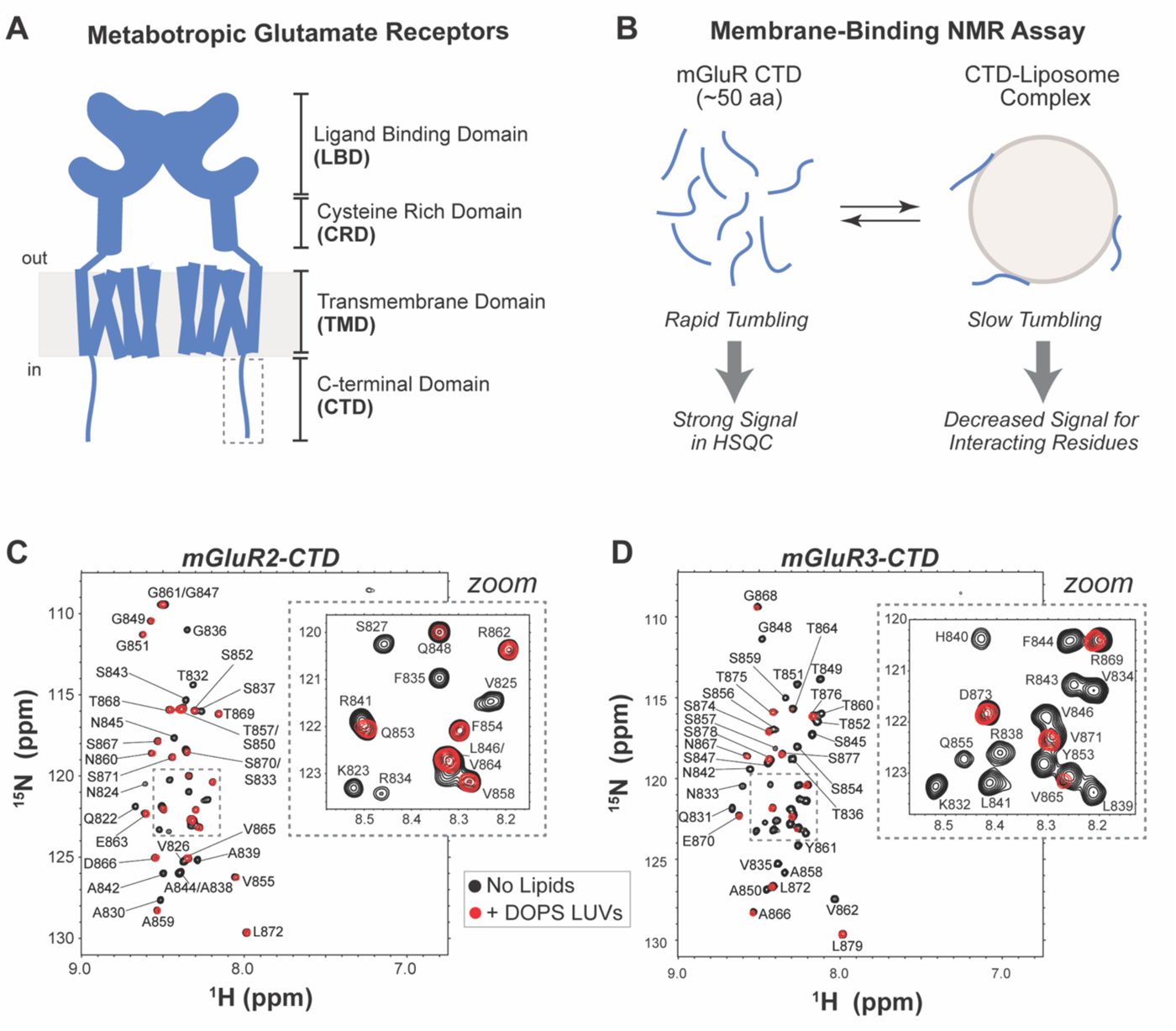
An NMR-based assay reveals phospholipid membrane binding of the intrinsically disordered mGluR2 and mGluR3 CTDs. **(A)** Schematic of the structural organization of mGluR domains highlighting the location of the CTD compared to the ordered parts of the protein and the membrane. **(B)** Schematic of NMR-based CTD-membrane binding assay that takes advantage of changes in tumbling rates of the CTD due to interactions with LUVs. **(C)** ^1^H-^15^N HSQC spectra of isolated mGluR2* and **(D)** mGluR3* CTD in the presence (red) and absence (black) of 10 mM 100 nm diameter LUVs comprised of DOPS at pH 6.8 at 10 °C, with zoomed insets of crowded regions, highlighting loss of signal of specific residues in the presence of LUVs.

The central role of the CTD in subtype-specific mGluR regulation raises the question of its structural properties and whether its proximity to the membrane may shape its structure or function. Despite recent cryo-EM studies which have resolved the structures of mGluR ECDs and TMDs in inactive and active states^9,29,35–38^, structural information on group II mGluR CTDs is limited to a short membrane-proximal segment of the mGluR2 CTD (residues 821 to 830) which was observed bound to G protein in a recent report^9^. Here, we examine the structural properties and membrane-interactions of the CTDs of mGluR2 and mGluR3. Using a combination of spectroscopic and computational approaches we find that both mGluR2 and mGluR3 CTDs are intrinsically disordered but are capable of interaction with phospholipid membranes. We show that both TMD-proximal and -distal basic residues can mediate electrostatic interactions with lipid headgroups. Additionally, we identify in mGluR3 an aromatic residue, Y853, that can partition into the membrane interface to potentially restrict access to the mGluR3 CTD S/T-rich. We then find that point mutations, including those associated with melanoma, can modulate membrane interactions, β-arr-dependent internalization, and G protein activation in cells. Furthermore, we show that EGF stimulation leads to agonist-independent mGluR3 internalization, depending on the presence of Y853, suggesting that Y853 phosphorylation may drive receptor internalization by reducing binding of the CTD to the membrane. Together this work suggests complex and dynamic interactions between the intrinsically disordered CTD and the membrane, expanding the known repertoire of GPCR regulatory mechanisms.

## Results

### Disordered mGluR CTDs bind negatively charged membranes *in vitro*

To investigate the structure of mGluR CTDs and probe potential membrane interactions, we turned to nuclear magnetic resonance (NMR) spectroscopy using purified, recombinant CTD constructs. For a quantitative, single-residue resolution assay of membrane binding, we took advantage of the effects of the slow tumbling rates of large unilamellar vesicles (LUVs) on NMR signals. Briefly, interactions of intrinsically disordered proteins (IDPs) with vesicles attenuate the signals of residues that interact with the membrane because they adopt the slow tumbling rates and long rotational correlation times of the LUVs (**Fig. 1B**). This assay has been used extensively to characterize IDP/membrane interactions^39–41^ but has not been applied to GPCR CTDs. We first obtained 2D ^15^N-^1^H HSQC spectra for both mGluR2 and mGluR3 CTDs in the absence of lipids, which exhibited the sharp signals and limited dispersion that are characteristic of IDPs (**Figs. 1C,D**). In the presence of LUVs composed of DOPS (18:1 1,2-dioleoyl-sn-glycero-3-phospho-L-serine) many signals in the spectra of both CTDs were clearly attenuated (**Figs. 1C,D insets**), indicative of an interaction between the corresponding CTD residues and the negatively charged vesicles. Spectra acquired in the presence of LUVs comprised of 1:1 DOPS:DOPC (18:1 [Δ9-Cis] 1,2-dioleoyl-sn-glycero-3-phosphocholine) or only DOPC exhibited variable degrees of signal loss (**Fig. S1**) suggesting that CTD-membrane interactions are sensitive to lipid composition.

We then asked whether the mGluR CTDs form helical secondary structure upon membrane binding using circular dichroism (CD) spectroscopy. CD spectra of the mGluR2 (**Fig. S2A**) and mGluR3 (**Fig. S2B**) CTDs showed no evidence for alpha-helix formation either in the absence of lipids or in the presence of vesicles under conditions where maximal membrane binding was observed by NMR. Because formation of short segments of helical structure can be difficult to detect in longer polypeptides, we also examined CD spectra of shorter mGluR2 and mGluR3 CTD peptides corresponding more closely to just their membrane-binding regions (see methods). These spectra (**Figs. S2C,D**) also showed no evidence of alpha-helix formation in the absence or presence of membranes. Even the presence of membrane mimetic SDS and DPC micelles, which often induce helical structure^42–44^, did not result in helix formation (**Figs. S2E**). As a control, we confirmed using CD that LUVs induce robust helical structure in a helix-8 peptide from NTS1 (**Fig. S2F**), as previously reported^45^. These results are consistent with the physicochemical characteristics of the membrane-binding regions of the mGluR2 and mGluR3 CTDs, which do not show the amphipathic nature typical of membrane-induced helices (see helical wheel plots in **Fig. S2G,H**) as well as with secondary structure predictions. Thus, it appears that the conformational ensemble sampled by group II mGluR CTDs upon membrane binding lacks persistent secondary structure and maintains a high degree of disorder.

### Electrostatics mediate mGluR CTD-membrane interactions

To enable sequence-specific analysis of CTD-membrane binding, we obtained NMR backbone resonance assignments using conventional triple resonance NMR experiments (**Figs. 1; S1**). Chemical shift-based secondary structure assessments confirmed the highly disordered nature of both CTDs in the absence of LUVs, as indicated by the lack of any significant secondary shifts (**Figs. S3A,B**) and corroborated using CheSPI^46^, which indicated negligible probabilities for helix or strand secondary structure. Plots of the ratio of NMR signal intensities in the presence versus absence of LUVs as a function of position (**Figs. 2A,B**) show that both mGluR CTDs interact with phospholipids via their N-terminal regions. For both CTDs, NMR signal attenuation is dependent on negatively charged lipid content, as signal attenuation is absent (mGluR2) or decreased (mGluR3) in the absence of DOPS (**Figs. 2A,B; S1**). Despite similar binding profiles at their N-termini, the membrane binding region for the mGluR3 CTD is longer, spanning the first ∼30 residues, compared with ∼21 residues for mGluR2. Plots of the average intensity ratio as a function of LUV composition within the N-terminal 20 residues (**Fig. 2C**) illustrate the clear DOPS-dependence of membrane binding in this region, in contrast with the C-terminal region (**Fig. 2D**).

**Figure 2:**
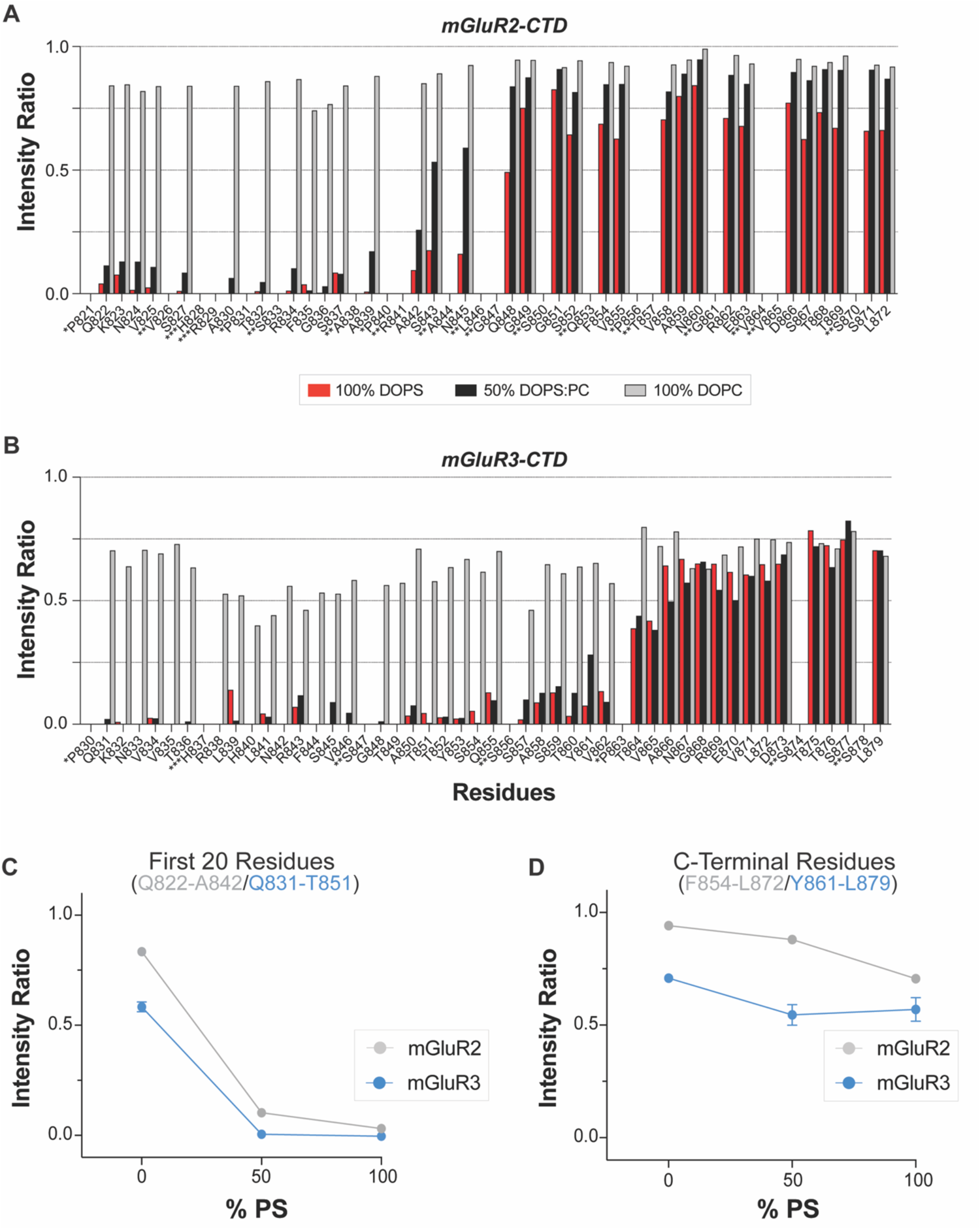
N-terminal regions of mGluR2 and 3 CTDs interact with negatively charged lipids. NMR intensity ratios for **(A)** mGluR2 and **(B)** mGluR3 from spectra collected with and without LUVs of three different lipid compositions. Prolines, which do not give rise to signals in ^1^H-^15^N HSQC spectra, are denoted by *, overlapping peaks for which values were not included by **, and residues not detected in the spectra by ***. **(C)** Averaged intensity ratios over the first ∼20 residues (mGluR2 Q822-A842; mGluR3 Q831-T851) illustrate the lipid composition dependence of the interactions in this region (±s.e.m.). **(D)** Averaged intensity ratios over the last ∼20 residues (mGluR2 Q853-L872; mGluR3 Y861-L879) illustrate the lack of lipid composition dependence of the interactions in this region (±s.e.m.).

Inspection of the sequence of both mGluR CTDs revealed a conserved N-terminal cluster of basic residues (**Fig. 3A**), which we hypothesized could drive CTD binding to negatively charged DOPS headgroups. Mutagenesis of each of the four basic residues in this cluster to alanine reduced membrane binding for this region of the mGluR2 CTD (**Figs. 3B-D; S4**), with the strongest effect observed for LUVs composed of 1:1 DOPS:DOPC, indicating that increasing negative charge content in the membrane can partly compensate for the loss of individual basic residues in the CTD (**Fig. 3C**). The strongest effect was observed for the R834A mutation, and mutation of the corresponding residue in mGluR3 to alanine (R843A) also lead to severe disruption of membrane binding by the mGluR3 CTD (**Fig. 3E**). These results support a major role for electrostatic interactions between basic CTD residues and anionic phospholipids in driving CTD-membrane interactions.

**Figure 3:**
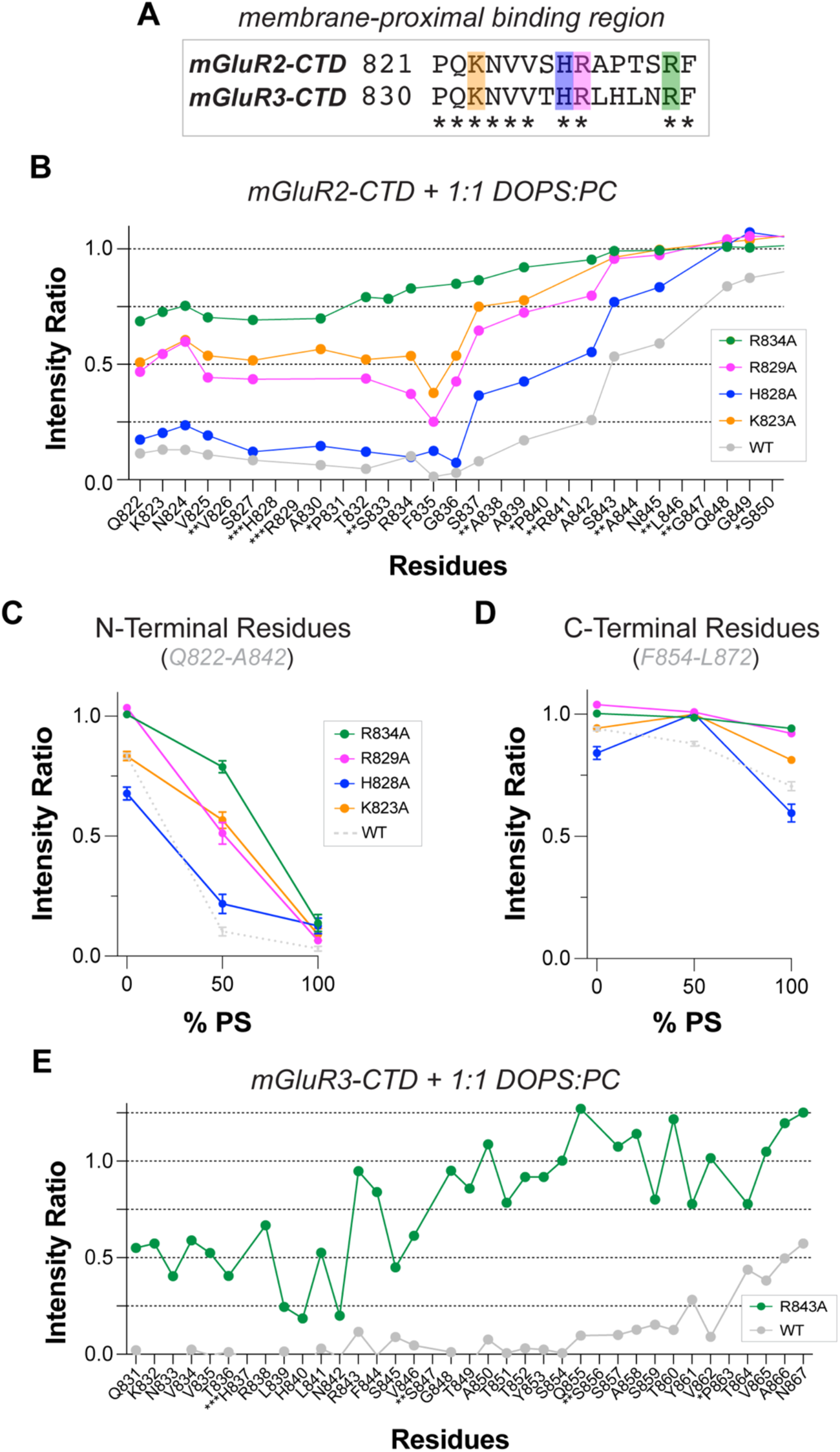
An N-terminal cluster of basic residues is critical for CTD membrane binding. **(A)** sequence alignment of the first 15 residues of the mGluR2 and mGluR3 CTDs highlighting conserved (*) and positively charged (highlighted) residues. **(B)** Intensity ratio plots of mGluR2 CTD constructs containing alanine substitutions for each of the four basic residues from spectra collected with and without LUVs containing a 1:1 mixture of DOPS:DOPC lipids. **(C)** Averaged intensity ratios over the first ∼20 residues (Q822-A842) illustrate the regional effect of each mutation for LUVs of different lipid composition (±s.e.m.). **(D)** Averaged intensity ratios over the last ∼20 residues (Q853-L872) illustrate the regional effect of each mutation for LUVs of different lipid composition (±s.e.m.). **(E)** Intensity ratio plots of R843A mGluR3 CTD compared to WT from spectra collected with and without LUVs containing a 1:1 mixture of DOPS:DOPC lipids.

To further understand the properties of the mGluR CTD-membrane interaction, we examined the potential role of membrane curvature in altering the membrane-bound region of mGluR CTD. Notably, disordered protein segments can sense membrane curvature on short length scales by interacting with lipid packing defects, without the need for shape complementarity over longer distances^47,48^. We measured binding to DOPS vesicles with diameters ranging from 50-400 nm and observed no substantial changes in mGluR2 membrane binding, indicating that mGluR CTDs are insensitive to membrane curvature (**Fig. S5**). To assess whether different regions of the protein bind to membranes with different affinities, we obtained NMR intensity ratio data for the mGluR3 CTD as a function of lipid concentration (**Fig. S6A**). We found that the N-terminal basic cluster remains bound even at lower lipid concentrations but observed a reduction in binding of the subsequent residues (S845 to T860) comprising the S/T-rich region of the mGluR3 CTD, suggesting that this region binds less strongly (**Fig. S6C**). To explore how CTD membrane interactions are affected by more complex lipid compositions that more closely resemble the plasma membrane inner leaflet, we measured mGluR3 CTD binding to DOPE-containing (1,2-dioleoyl-sn-glycero-3-phosphoethanolamine) vesicles composed of 11:4:5 DOPC:DOPE:DOPS with or without 30% cholesterol. The 25% negative charge content of these vesicles is similar to that expected for the inner leaflet of cellular plasma membranes^49,50^. The resulting binding profile (**Fig. S6A)** exhibits strong binding in the basic cluster region, but attenuated binding in the S/T-rich region when compared with 1:1 DOPC:DOPS vesicles, consistent with reduced binding at lower negative charge content. Indeed, the profile closely resembles that observed in the presence of lower (2.5 mM) concentrations of 1:1 DOPC:DOPS vesicles (**Fig. S6A)**. The inclusion of 30% cholesterol did not alter the binding profiles, suggesting that cholesterol may not strongly influence CTD-membrane interactions (**Fig. S6A**).

Finally, to assess CTD membrane binding in a concentration-independent manner and in a context more closely resembling that of the intact membrane-inserted receptor, we anchored the mGluR3 CTD to vesicles by introducing a hexa-histidine tag at its N-terminus and doping 5% DGS-Ni-NTA (1,2-dioleoyl-sn-glycero-3-[(N-(5-amino-1-carboxypentyl)iminodiacetic acid)succinyl] (nickel salt)) into PC:PE:PS vesicles (10:4:5:1 DOPC:DOPE:DOPS:DGS-Ni-NTA). Intensity ratio plots for the anchored CTD revealed a signal attenuation profile consistent with that observed for the free CTD, with the strongest binding at the N-terminus and clear, but decreasing binding through the S/T-rich and C-terminal regions (**Fig. S6E**). Importantly, the extent of attenuation for the anchored CTD is dramatically greater in the S/T-rich and C-terminal regions than that observed for the unanchored CTD using vesicles containing 50% negative charge content. Because nickel is paramagnetic, DGS-Ni-NTA will also induce some degree of paramagnetic relaxation enhancement (PRE) of CTD nuclei that approach within a ∼25 Å distance, adding a distance-based signal attenuation to that resulting from immobilization of residues on the vesicle. To assess the extent of this effect, we measured intensity ratios for the unanchored CTD, lacking the N-terminal hexa-histidine tag, in the presence of DGS-Ni-NTA vesicles. This condition produced only minor broadening beyond that observed for corresponding vesicles lacking DGS-Ni-NTA, suggesting that NMR resonance attenuation induced by slow tumbling upon membrane binding dominates any DGS-Ni-NTA-associated PRE effects. Together, these results demonstrate that N-terminal anchoring of the mGluR3 CTD to vesicles results in enhanced membrane interactions, and indicate that the lipid concentrations and compositions and the sample conditions used in our *in vitro* studies of untethered CTD peptides do not result in artifactual binding profiles or overestimates of binding.

### Aromatic and distal charged residues help anchor the mGluR3 CTD S/T-rich region to the membrane

To further probe structural changes in the mGluR CTDs upon binding to membranes, we turned to all-atom molecular dynamics simulations. We focused our analysis on the mGluR3 CTD, as this subtype features more extensive membrane interactions than mGluR2 (**Figs. 2**), and since the CTD is known to have a central role in mGluR3 regulation^34^. We built a system containing a 1:1 DOPC/DOPS phospholipid bilayer and a protein chain comprising both transmembrane helix 7 (TM7) of mGluR3 and the CTD (*see Methods*). We included the transmembrane tether to increase the probability of observing CTD-membrane interactions within the simulation time and to bridge the *in vitro* experiments with isolated CTDs to the biological context of the CTD where it is attached to the TMD at its N-terminus. We started the simulations with the CTD in a disordered conformation with no contacts with the membrane and 6 independent replicas were simulated for 1,370 ns each, resulting in 8.22 µs of total simulation time (**Figs. S7A-F**, panels i). We did not observe any alpha-helix formation and found instead that the mGluR3 CTD is conformationally dynamic with little or no regular secondary structure (**Figs. S7A-F,** panels ii), consistent with our CD data (**Fig. S2**).

Our simulations consistently revealed conformations in which segments of the CTD were in contact with the membrane, especially in the N-terminal basic cluster region, where specific interactions between arginine residues and lipid headgroups were captured (**Fig. 4A; Movies SM1,2**). To quantify such interactions, we analyzed hydrogen bonding between CTD sidechains and lipid headgroups over all of our simulations. We found that H-bonds between basic cluster arginine residues (R838 and R843) and PS headgroups anchor the N-terminal region to the membrane and that R869 also forms such H-bonds (**Figs. 4B; S8A**). To more generally assess CTD-membrane interactions on a residue-by-residue basis, we calculated the distance of each sidechain in the CTD from the lipid phosphate plane. We find that when averaged over the individual replicas (**Fig. S7A-F**, panels iii) or over all the replicas (**Fig. 4C**) the N-terminal basic cluster region exhibits close proximity (<= 10 Å) to the membrane surface, consistent with the tighter binding observed for this region in our NMR experiments. Time courses of arginine sidechain-phosphate plane distances for different trajectory segments reveal that R838 and R843 exhibit long perios of close proximity to the membrane (**Fig. S8B,C**), whereas R869 exhibits more dynamic behavior, suggesting that it mediates transient anchoring of the C-terminal portion of the CTD to the bilayer.

**Figure 4:**
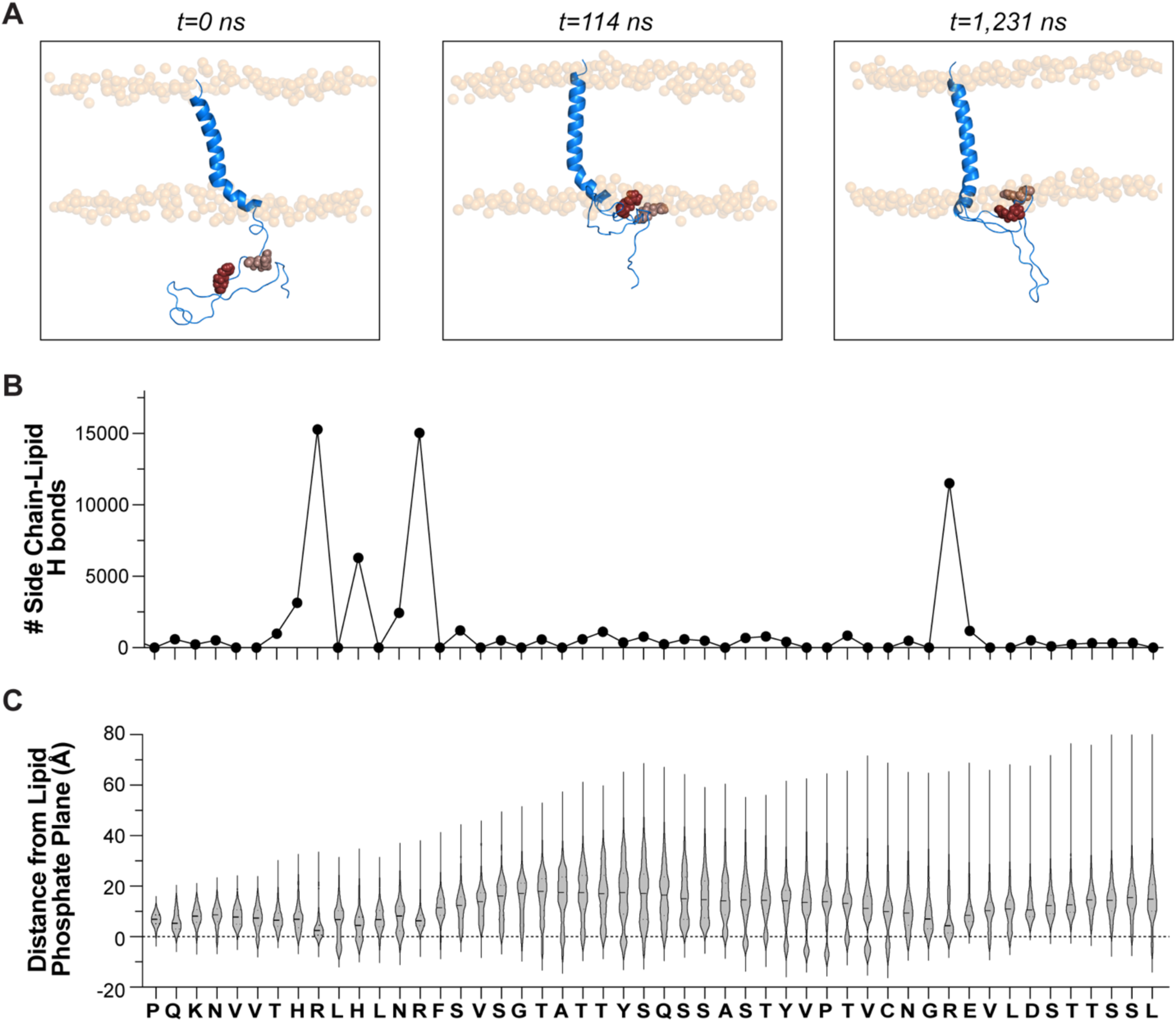
Molecular dynamics simulations reveal multi-modal membrane interactions of the intrinsically disordered mGluR3-CTD. **(A)** Snapshots of mGluR3 TM7-CTD (comprising TM7 residues 796-821 shown in cartoon helix representation, and CTD residues 822-879) replica 6 trajectory, highlighting three conformations of the CTD: the beginning of the simulation with no membrane contacts (t=0, left), and two membrane-associated states (t=174.4 ns (middle) and t=1,231 ns (right)). Protein backbone is in blue cartoon; R838 (brown) and R843 (red) sidechains shown as spheres) **(B)** Total number of hydrogen bonds between each side chain and lipid headgroups averaged over all simulations. **(C)** Diistributions, in the form of violin plots, of the distance of each residue (side chain center of mass) from the lipid phosphate plane over the course of all simulations (mean and quartiles depicted by solid and dotted horizontal lines, respectively).

For the S/T-rich region, the distance distributions appear to be bimodal, with a minor population that is very close to the membrane surface and a larger population that is more distant. The membrane-proximal states may represent a bound population, which appears smaller than that observed by NMR for the anchored CTD peptide (**Fig. S6E**), but unfortunately differences in the experimental and simulation conditions likely preclude a direct comparison. More generally, the simulations are consistent with the S/T-rich region binding the membrane less tightly than the N-terminal basic cluster region, as indicated by our NMR data. Intriguingly, residue Y853 within the the S/T-rich region (**Fig**. **5A**), which we have shown confers β-arr-mediated internalization of mGluR3^34^, exhibits a population with negative distances from the lipid phosphate plane, indicating that its side chain inserts at least partially into the membrane. Aromatic residues are known to partition favorably into membranes and are often found in interfacial regions of transmembrane domains^51–53^. We examined the time course of the phosphate plane distance of Y853 and identified long (400-650 µs) time periods in separate replicas during which this distance was negative or very small (**Fig. 5B,C; Fig. S8D**). Individual poses of Y853 showed its sidechain inserted into the membrane or at the interfacial lipid headgroup region (**Figs. 5B; Movies SM1,2**). Furthermore, the membrane proximity of nearby S/T residues appears to be correlated with that of Y853 (**Fig. S8E**), while remaining dynamic (**Fig. S8F**). To further probe the role of this aromatic residue in the membrane interactions of the S/T-rich region of the mGluR3 CTD, we examined the effects of mutating Y853 to alanine using our NMR-based assay. Compared to WT, Y853A resulted in a similar binding profile to 1:1 DOPC:DOPS vesicles for the N-terminal basic region, but showed decreased binding of the CTD in the S/T-rich region (**Figs. 5D; S9A,B**). Indeed, the resulting profiles resemble those obtained at lower lipid concentrations (**Fig. S6A**), indicating that Y853 helps to stabilize the membrane bound state of the S/T-rich region that we observe at higher lipid concentrations.

**Figure 5:**
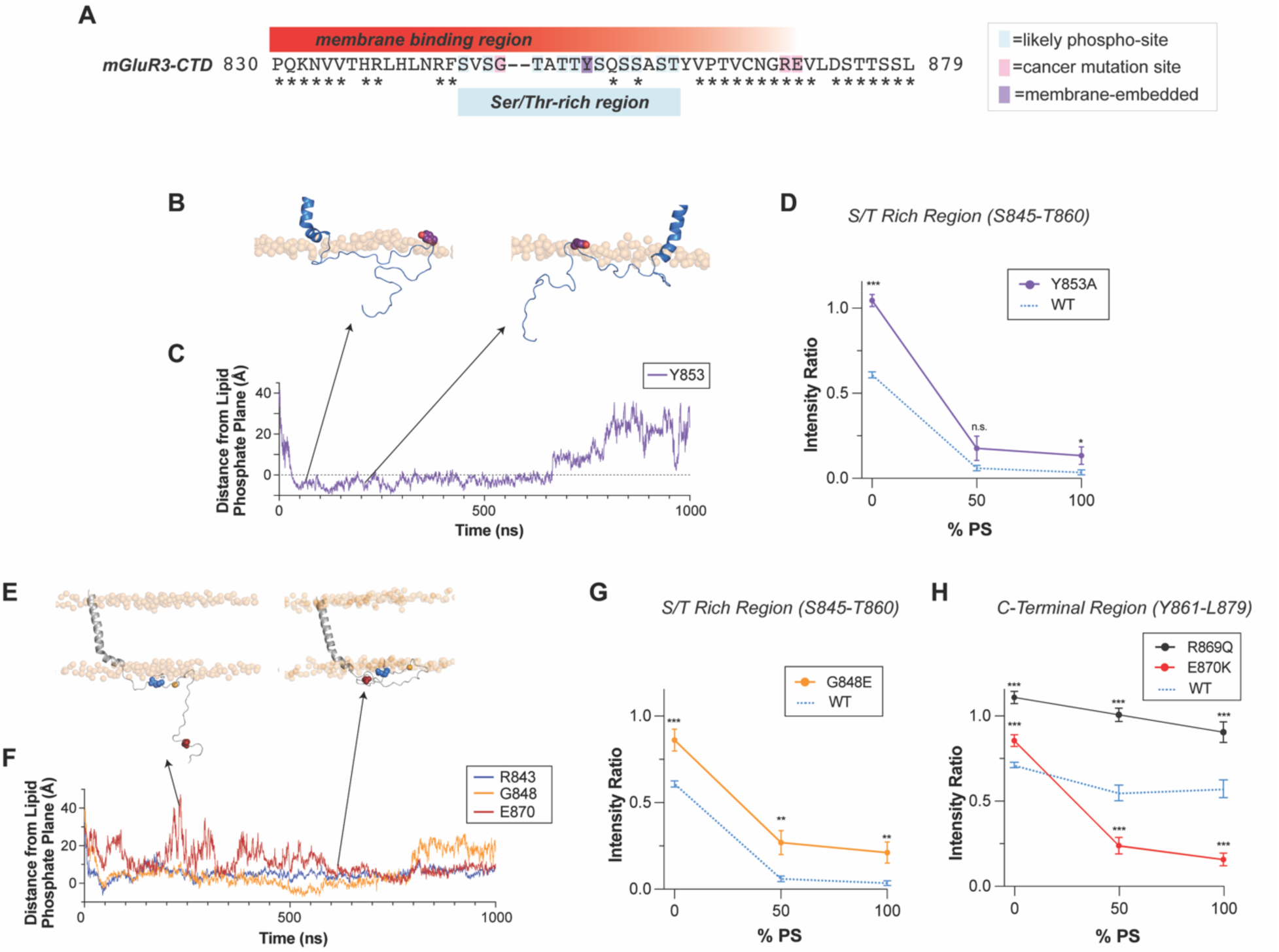
Membrane interactions of the S/T-rich region of the mGluR3 CTD is modulated by mutation of a key residue and by cancer mutations. **(A)** mGluR3-CTD sequence annotated with the NMR-and MD-determined membrane binding region and the overlapping Ser/Thr-rich region (* denotes residues conserved in mGluR2). **(B)** Snapshots of residue Y853 (shown as violet spheres with the hydroxyl group in red) in membrane-embedded and membrane-associated positions (from MD replica 6). Lipid phosphates are shown as transparent orange spheres. **(C)** Distance of Y853 sidechain to the lipid phosphate plane plotted as a function of time for MD replica 6 (first 1,000 ns). **(D)** Comparison of the averaged integrated NMR intensity ratios of WT mGluR3-CTD (dotted blue) with Y853A mGluR3-CTD taken over the S/T-rich region (S845-T860) as a function of LUV lipid composition (±s.e.m.; Wilcoxon test; *p<0.05, ***p<0.001, n.s. p≥0.05). **(E)** Snapshots of residues R843 (blue), G848 (orange) and E870 (red) at different time points during the time course of MD replica 6 showing prolonged membrane-association of R843 and G848 and fluctuating membrane-association of E870 (protein backbone is in gray cartoon; side chains shown as spheres colored as in **(F)** below; lipid phosphates are shown as transparent orange spheres). **(F)** Position of side chains R843, G848, and E870 relative to the phosphate plane of the membrane (see methods) throughout the time course of MD replica 6. **(G)** Comparison of the averaged NMR intensity ratios of WT (dotted blue) with G848E (orange) mGluR3-CTD taken over the S/T-rich region (S845-T860) as a function of LUV lipid composition (±s.e.m.; Wilcoxon test; **p<0.01, ***p<0.001). **(H)** Comparison of the averaged NMR intensity ratios of WT (dotted blue) with R869Q (black) and E870K (red) mGluR3-CTD taken over the last 19 residues (Y861-L879) as a function of LUV lipid composition (±s.e.m.; Wilcoxon test; ***p<0.001).

Interestingly, the distance distributions of the C-terminal region of the CTD indicates that a hydrophobic region preceding residue R869 also features a population with negative distances from the membrane plane (**Fig. 4C**; residues Y861-C866). Although this is not strikingly evident in the NMR data, for both the unanchored (**Fig. 2B**) and anchored (**Fig. S6E**) CTD, we observe a dip in the NMR intensity ratios in this region, and the NMR data for the anchored CTD show significant binding for the entire C-terminal region.

Noting that membrane-proximal conformations tended to be more extended in our simulations (**Fig. 5B**) we considered whether the radius of gyration (R_g_) of CTD conformations correlated with membrane proximity. We observed a bimodal distribution of R_g_ in our simulations, with a pinch point around the average R_g_ value of 15.5 Å (**Fig. S8G**). We calculated the average distance from the membrane for the ensemble of conformations with R_g_ below or above 15.5 Å and observed that conformations with lower R_g_ were biased towards shorter distances from the membrane (**Fig. S8H**). We extended this analysis by separately considering, as a function of R_g_, the distance of the N-terminal basic cluster region, the S/T-rich region and the C-terminal region from the membrane. For the N-terminal region, the distance from the membrane was small (<= 10 Å) irrespective of R_g_, consistent with tight binding of this region to the membrane. For the S/T-rich and C-terminal regions, compact conformations were distributed closer to the membrane, while more extended conformations were distributed further from the membrane, consistent with more dynamic and reversible interactions with the membrane. These results, which can also be appreciated in **Movies SM1,2**, suggest that membrane-binding restricts the conformational space of the CTD. Notably, both the S/T-rich and C-terminal regions featured a cluster of compact conformations situated very near (<= 10 Å) the membrane surface, consistent with the sidechain-phosphate plane distance distributions (**Fig. 4C**).

### CTD-membrane binding is modulated by cancer-associated mutations

The mGluR3 CTD contains a number of cancer-associated mutations, two of which, G848E and E870K, are associated with melanomas^54^ and a third, R869Q, which is enriched in carcinomas. While the role of these mutations in cancer remains unclear, each has been identified in multiple samples of cancer tissues (5, 4 and 6 samples for G848E, R869Q and E870K) according to the COSMIC database^55^. Multiple occurrences of identical mutations are statistically unlikely (estimated at 2E-12 for E870K^54^) and E870K has been also shown to increase melanoma cell growth, migration, and metastasis^54^. Interestingly, each of these mutations alters the charge of the mGluR3 CTD, suggesting they could influence membrane interactions. G848 lies within the S/T-rich region, situated between R843 and Y853, and is membrane-associated according to both the NMR data and our MD simulations (**Figs. 2B, 4C; 5E,F; Movies SM3,4**). R869 and E870 are in the C-terminal region of the mGluR3 CTD that is more weakly membrane associated according to the NMR data and features transient contacts with the membrane in our simulations (**Figs. 2B, 5E,F; Movies SM3,4**). Based on these results, we hypothesized that these cancer-associated mutations could alter membrane binding due to changes in the local electrostatic properties of the mGluR3 CTD that would either diminish (G848E, R869Q) or promote (E870K) interactions with DOPS headgroups. To test this, we measured binding of these mutants using our NMR-based approach. The mGluR3 CTD G848E variant resulted in reduced interaction between the S/T-rich region of the CTD and the membrane, similar to the effect observed for the Y853A mutation (**Figs. 5G; S9A,B**). Strikingly, the R869Q mutation dramatically decreased membrane binding of both the S/T-rich region and of the C-terminal region of the CTD (**Figs. 5H; S9A,B**). In contrast, the E870K mutation extended the membrane-interacting region of the CTD nearly to its very C-terminus (**Figs. 5H; S9A,B**). The ability of these cancer-associated mutations to alter CTD-membrane binding further supports the role of electrostatics in driving these interactions.

### mGluR3 CTD membrane interactions modulate receptor internalization in living cells

Having established that mGluR3 mutations can alter membrane binding *in vitro*, we next asked if modifications which alter CTD-membrane interactions can also alter receptor function in living cells. We initially focused on agonist-induced mGluR3 internalization, which is driven by phosphorylation-dependent interactions with β-arrs^34^. As described above, we reasoned that binding of the mGluR3 S/T-rich region to the membrane surface could modulate the ability of the CTD to interact with GRKs and/or β-arrs to mediate internalization.

We assessed the effects of the R843A, G848E, Y853A, R869Q and E870K mutations (**Fig. 6A**) on mGluR3 internalization using an established live cell surface labeling imaging-based assay^34^. All point mutants expressed on the surface, although a small decrease relative to wild type was observed for G848E (**Fig. S10A**). To quantify receptor internalization, we labeled N-terminal SNAP-tagged mGluR3 transfected into HEK 293T cells with a membrane-impermeable fluorophore after 60 min treatment with agonist or antagonist. A consistent ∼30% drop in fluorescence, reflecting receptor internalization, was observed for wild type mGluR3 following agonist treatment, reflecting endocytosis (**Fig. 6B**). Compared to WT, the R843A, G848E, Y853A, and R869Q mGluR3 mutants exhibit a greater degree of glutamate-evoked internalization (**Fig. 6B**). The data for G848E are consistent with our previous report that this mutation results in enhanced internalization^34^. In contrast, the E870K mutation drastically decreased glutamate-induced internalization of mGluR3 (**Fig. 6B**). These results are consistent with our hypothesis that CTD-membrane interactions regulate the accessibility of the CTD to GRKs and β-arrs, with mutations that inhibit or enhance CTD-membrane binding exhibiting enhanced or blunted internalization, respectively.

**Figure 6:**
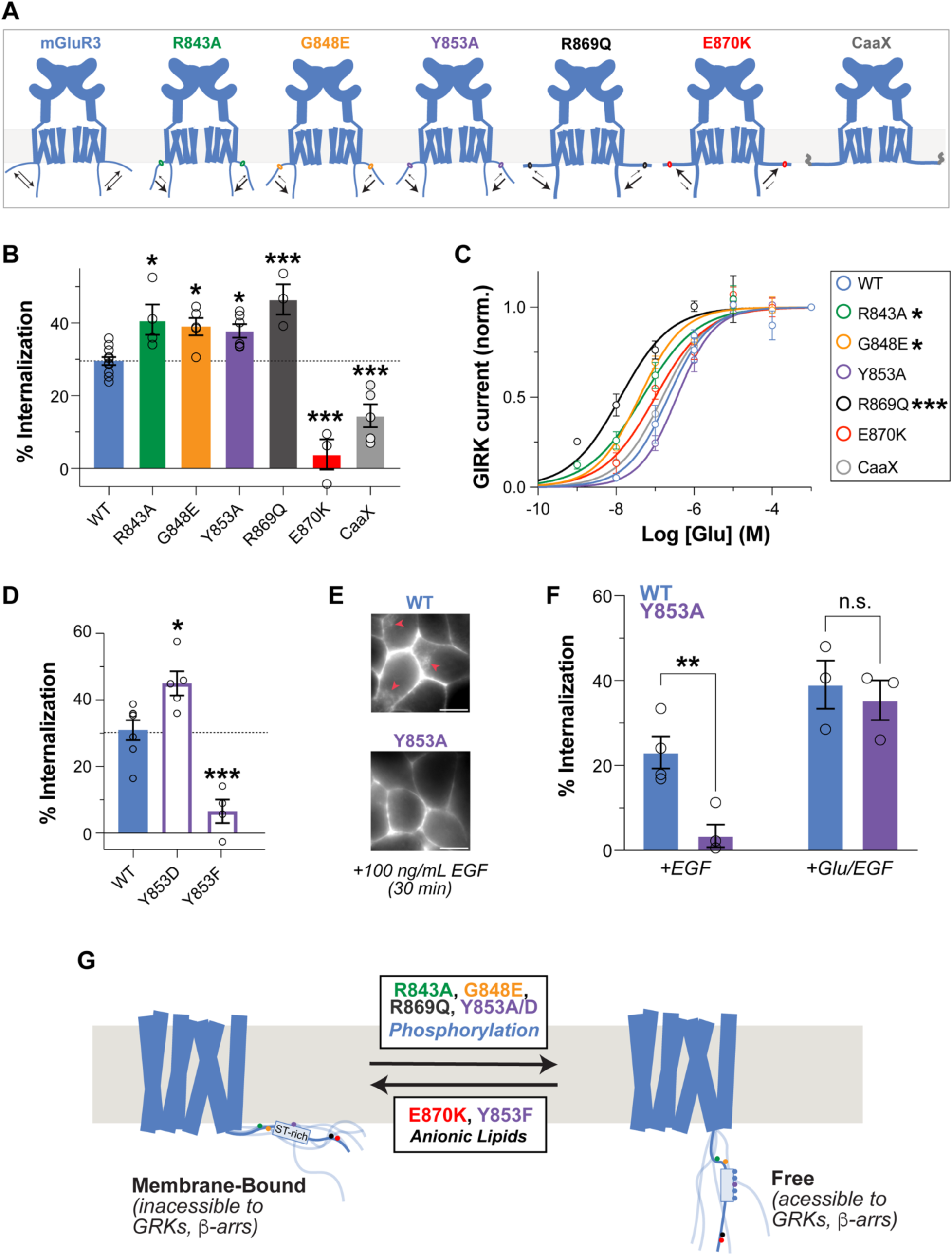
CTD mutations that alter membrane binding affect mGluR3 internalization and function. **(A)** Schematics of mGluR3 CTD mutational positions and their effects on mGluR3-CTD free vs. membrane-bound equilibrium. Larger arrows show the direction in which each variant perturbs the equilibrium. **(B)** Quantification of the extent of receptor internalization for each mGluR3 variant (with dotted line denoting mGluR3 WT internalization) (averaged internalization per day, 10-12 images per condition/day and 4-9 days per condition; One-Way ANOVA with multiple comparisons, * p<0.05 *** p<0.001). **(C)** Glutamate dose response curves for mGluR3 variants in a patch-clamp experiment using GIRK currents as a reporter for mGluR3 G-protein activation (EC50: WT = 136.9 ± 26.9 nM, R843A = 51.2 ± 11.9 nM, G848E = 44.2 ± 8.6 nM, Y853A = 417.8 ± 64.8 nM, R869Q = 14.1 ± 4.3 nM, E870K = 101.6 ± 21.7 nM, CAAX = 156.9 ± 42.8 nM; F-test of EC50 shifts; **p<0.01, ***p<0.001). **(D)** Quantification of the extent of receptor internalization of WT mGluR3 vs. Y853D phospho-mimetic mutant vs. Y853F (averaged internalization of 10 images per condition/day across 3 days; t-test; *p<0.05). **(E)** Representative images of HEK293T cells expressing SNAP-tagged mGluR3 WT vs Y853A treated with 100 ng/ml EGF for 30 min (red arrows represent internalization; scale bar: 5 µm). **(F)** Quantification of the extent of internalization for mGluR3 WT vs Y853A mutant in EGF or Glu+EGF incubated conditions (averaged internalization per day, 10 images condition/day and 3-4 days per condition; t-test, **p<0.01, n.s. p≥0.05). **(G)** Working model of mGluR3-CTD free vs. membrane-bound equilibrium and changes that favor the less accessible membrane-bound (E870K, Y853F, anionic lipids) or the more accessible free (R843A, G848E, R869Q, Y853A/D, phosphorylation) state.

To further assess our interpretation that altered CTD-membrane interactions underlie the observed changes in receptor internalization, we examined the effects of artificially anchoring the mGluR3 CTD to the membrane by appending a CAAX box lipidation motif to its C-terminus (**Figs. 6A; S10A**). Indeed, this variant exhibited a reduction in glutamate-induced receptor internalization similarly to the E870K mutant receptor (**Fig. 6B**). We also visualized receptor internalization via live cell microscopy where we labeled plasma membrane SNAP-tagged mGluR3 variants with a membrane-impermeable fluorophore and visualized fluorescence localization following glutamate treatment (30 min, 1 mM). This analysis confirmed the enhanced internalization of R843A, G848E, Y853A and R869Q and the reduced internalization of E870K and mGluR3-CAAX following glutamate treatment (**Fig. S10B**).

To assess any effects of CTD mutations on G protein activation, we performed patch clamp measurements using a GIRK channel current assay (see Methods). In this assay, R843A, G848E and R869Q showed a clear left-shift while Y853A, E870K and -CAAX did not show significantly different apparent glutamate affinities compared to wild type mGluR3 (**Figs. 6C; S10C,D**). These observations suggest that mutations can exert distinct functional effects on β-arr and G protein coupling.

Motivated by our observations of the contributions of residue Y853 to the membrane interactions of the mGluR3 CTD S/T-rich region, we posited that phosphorylation of this residue could influence CTD-effector interactions by reducing membrane binding. Consistent with this hypothesis, a phosphomimetic Y853D mutant showed enhanced internalization compared to WT (**Fig. 6D**) and also resulted in dramatically decreased membrane binding of the S/T-rich region (**Fig. S10E**). We then found that treatment of mGluR3-transfected HEK 293T cells with epidermal growth factor (EGF; 100 ng/ml), which stimulates myriad downstream kinase signaling pathways, led to detectable internalization of WT, but not of Y853A mGluR3 in the absence of agonist (**Figs. 6E,F**). Combining agonist and EGF treatment enhanced internalization to a similar extent to that observed for the agonist-treated Y853A mutant (**Fig. 6B**). Interestingly, the addition of EGFR tyrosine kinase inhibitor AG1478 eliminated EGF-induced internalization, while the application of Dasatinib, a pan-Src family tyrosine kinase inhibitor, did not produce a significant change in the EGF-induced internalization compared to the control (**Fig. S10H**). This points towards a direct effect of the EGFR tyrosine kinase domain in either directly phosphorylating Y853 or activating a different downstream pathway that, ultimately, targets mGluR3. Together these results suggest that membrane-interacting residues in the mGluR3 CTD can contribute to both agonist-driven homologous internalization and heterologous internalization following stimulation of other cellular pathways.

In principle, the enhancement of mGluR3 internalization by Y853D could reflect increased binding to β-arr caused by mimicking phosphorylation within the S/T-rich region. The fact that Y853A also enhances internalization argues against this possibility, since it is unclear how that replacement of Y853 with an alanine would promote binding to β-arr. Nevertheless, we explored a subtler change at this position by replacing Y853 with phenylalanine. This mutation, which removes only one hydroxyl group, would not be expected to have a dramatic effect on any interaction with β-arr, but would be expected to enhance membrane binding by removing the polar hydroxyl group that restricts Y853 to the membrane interface region. Accordingly, we found that the Y853F mutation dramatically decreases receptor internalization (**Fig.6D**). To verify the expected effect of this mutation on membrane binding, we examined binding at a series of lipid concentrations and showed that Y853F maintains strong binding of the S/T-rich region even at lower lipid concentrations, where binding by the WT CTD is decreased (**Fig. S6B,D**).

## Discussion

The functional roles of disordered intracellular domains in GPCRs, particularly their CTDs, have drawn increasing interest in recent years. Several studies have confirmed the disordered nature of GPCR CTDs^3–6^ and direct interactions of receptor CTDs with the intracellular face of the corresponding TMDs, which are regulated by phosphorylation and/or agonist binding and influence both receptor activity and coupling to β-arrs, have been documented^5,6,56^. Direct membrane binding of disordered intracellular domains has been shown to play functional roles for a number of membrane proteins ^57–59^, but GPCRs have not been the subject of such studies to date. Membrane phospholipid composition and cholesterol levels have been shown to modulate GPCR function^21,22^, but this has been thought to occur primarily by direct interactions with membrane-embedded TMDs. Here, we show that family C GPCRs mGluR2 and mGluR3 can also sense the membrane in a functionally relevant way through their disordered intracellular CTDs, and that modulating CTD-membrane interactions alters receptor internalization.

We recently reported that mGluR3, but not mGluR2, couples strongly to β-arrs, dependent on the presence of an S/T-rich region in its CTD^34,60^. Here we show that both CTDs are highly disordered in solution and bind to unilamellar lipid vesicles via their N-terminal regions. While many GPCRs feature a short amphipathic membrane-associated helix-8^45^, the mGluR2 and mGluR3 CTDs do not form detectable helical structure upon membrane binding. This is consistent with recently reported cryo-EM structures of full-length mGluR2 and mGluR3 which do not feature a classical helix 8^9,38,61^. Notably, our data do not rule out the possibility of very short helical segments in the membrane-bound CTDs, which have been observed in other intracellular membrane-binding domains^57^.

We note precedents for IDP-membrane interaction modes that do not involve secondary structure formation, including the MARCKS-ED peptide from the effector domain of myristoylated alanine-rich C-kinase substrate^62,63^, which features 13 positively charged and 5 hydrophobic residues within a short 25-residue polypeptide segment. The C-terminal motif of worm complexin also binds to membranes without secondary structure via a combination of positively charged and hydrophobic side chains^47,64^. Another particularly relevant example is the N-terminal region of the *Mycobacterium tuberculosis* divisome protein ChiZ, which binds to acidic membranes primarily via hydrogen bonds between phospholipid headgroups and 9 arginine residues^65^.

We posit that CTD-membrane interactions can regulate CTD availability for interactions with downstream effectors such as GRKs and β-arrs (**Fig. 6G**). We demonstrated that mutations that reduce membrane association (R843A, G848E, Y853A, Y853D and R869Q) result in increased receptor internalization, whereas mutations or modifications that enhance membrane binding (Y853F, E870K and introduction of a CAAX box motif) result in decreased receptor internalization. Importantly, half of these modifications are distant from the S/T-rich region of mGluR3 and are therefore unlikely to directly impact binding to GRKs or β-arrs. The presence of Y853 within the S/T-rich region and its importance for the membrane interactions of this region prompted us to hypothesize that phosphorylation of this tyrosine residue could also modulate CTD-membrane binding and thereby regulate coupling to transducers, including β-arrs. Notably, tyrosine phosphorylation has been reported to disrupt localized membrane binding of several disordered proteins^66,67^. While not providing direct proof, the dependence of EGF-induced mGluR3 internalization on the presence of Y853 and its elimination by an EGF kinase inhibitor support this possibility, as does the enhanced internalization we observe for the phosphomimetic Y853D mutation. This would also be consistent with literature reports of heterologous receptor internalization/desensitization^68^. Alternative explanations may exist and could include EGFR activation of GRK2 and downstream Ser/Thr phosphorylation in the CTD^69^, but it is unclear how this could account for the requirement for Y853. In light of our observations and of previous reports of the role of tyrosine-membrane interactions in regulating T cell receptor activation^57^, we posit that modulation of such interactions, either directly by phosphorylation or indirectly by other mechanisms may constitute a general mechanism for receptor regulation.

The work presented here makes a case for a general role for GPCR CTD-membrane interactions in regulating the accessibility of receptor CTDs to downstream effectors, including β-arrs (**Fig. 6G**). An appealing aspect of this model is that it provides a mechanism for sequential or cooperative phosphorylation^70^ as initial phosphorylation events could shift the equilibrium of phosphocode-containing regions^14,71^ and increase their accessibility for further phosphorylation and subsequent β-arr binding. Together with recent results describing functionally important interactions of receptor CTDs with the intracellular face of their TMDs^5,6,56^, our work expands the modalities by which GPCR CTDs can regulate receptor function. Important questions remain regarding how the interplay of CTD interactions with membranes, TMDs, G-proteins, GRKs, β-arrs and other effectors is regulated and orchestrated. Mutations that alter CTD-membrane interactions could also affect direct CTD G-protein binding/recruitment^7–9^, autoinhibitory CTD interactions with G-protein binding sites on the TMD^5,34,56^, or allosteric effect on TMD conformation, especially since these other interactions may also include electrostatic components. Indeed, we observe that mutations that strongly disrupt CTD-membrane binding facilitate receptor activation and we also recently reported an auto-inhibitory effect of the mGluR2 CTD on β-arr coupling^34^, supporting a potential interplay between G-protein-, TMD-and membrane-binding. Recent studies of the calcium sensing receptor (CaSR) also proposed a potential interplay between sequestration of a short CTD segment and its interactions with G-proteins^72^. While our focus here has been on the CTD, similar mechanisms and interactions may be operative for other disordered intracellular GPCR domains^73^.

## Conclusions

Our results establish a previously unappreciated yet critical and dynamic role of CTD membrane interactions in controlling GPCR desensitization and internalization and suggest that an equilibrium between membrane-bound and free states controls transducer coupling efficiency. This equilibrium may be modified in multiple ways, including disease mutations, Ser/Thr phosphorylation and possibly Tyr phosphorylation, as well as changes in membrane composition, comprising a novel mode of CTD-mediated GPCR regulation.

## Methods

Recombinant proteins were expressed in bacteria as fusion proteins and purified by affinity chromatography. Large unilamellar vesicals were prepared from the appropriate lipid composition by extrustion. NMR 2D experiments were collected at 500 MHz and triple-resonance experiments for assignments were collected at 800 MHz. Circular dichroism measurments were performed on an AVIV 410 CD spectropolarimeter over a wavelength range from 300-190 nm. MD simulations of an mGluR3 construct containing both TM7 and the CTD (residues 796-879) used initial poses generated using AlphaFold2^75^ and ColabFold^76^ which were equilibrated using the standard CHARMM-GUI based protocol and scripts followed by a short, 6-ns run using OpenMM^85^ and the CHARMM36m^86^ forcefield and then simulated for 1,370 ns for each of 6 replicas. Whole-cell patch clamp recordings were performed in HEK 293 cells 24 hours post-transfection as previously described^92^. For quantifying receptor internalization we used a previously reported surface labelling assay^34^. Details of all methodologies applied in this study are included in the SI Appendix.

## Data Sharing Plan

All NMR chemical shift assignments can be obtained online from the biological magnetic resonance database (BMRB accession numbers 52206 and 52202, to be released upon publication). NMR intensity ratio data, CD data, MD trajectories and all code used for the analysis of MD simulations can be obtained online at GitHub (https://github.com/cmanci/mGluR_CTD). Imaging data will be made available upon request, as well all plasmids and reagents used in the study.

## Acknowledgments and funding sources

This work was funded in part by National Institutes of Health (NIH) grants R35GM136686 (DE), R01NS129904 (JL), F32GM148001 (DM), F31AG077836 (CM) and the Margarita Salas Fellowship from the Ministry of Universities of Spain (A.G.-H), the Rohr Family Research Scholar Award (JL), and the Monique Weill-Caulier Award (JL). G.K. gratefully acknowledges support from the 1923 Fund. The WCM NMR core is supported by NIH shared instrumentation grants S10OD028556 and S10OD016320. We acknowledge assistance from Clay Bracken and Emily Grasso with NMR data collection and processing, Jihye Kim for guidance on EGF stimulation studies and Derek Shore for assistance setting up simulations. The authors gratefully acknowledge the use of in-house computational resources of the David A. Cofrin Center for Biomedical Information in the Institute for Computational Biomedicine at Weill Cornell Medical College.

## SI Appendix

### Protein expression and purification

Plasmids encoding the rat mGluR2 (residues 821-872) and mGluR3 (residues 830-879) CTDs preceded by an N-terminal 6x-His-SUMO tag were procured from Twist Biosciences. Cysteine-free versions of both mGluR2 (C859A) and mGluR3 (C866A) were generated using an In-Fusion Cloning kit (Takara Bio) and confirmed by DNA sequencing, and were used as the background for generating the other mutants described. In each case, a conserved proline located at the end of TM7^29,74^ was selected as the start of the CTD. Recombinant proteins were expressed in *E. coli* BL21/DE3 cells (Novagen) grown in either LB Broth or M9 minimal media containing ^15^N-labeled ammonium chloride (1 g/L) or ^15^N-labeled ammonium chloride and ^13^C-labeled D-glucose (2 g/L) at 37 °C (275 rpm) induced with 1 mM IPTG (Isopropyl β-D-1-thiocalactopyranoside) at OD6_00nm_ of 0.6-0.8. 4 hours post induction, cells were harvested via centrifugation at ca. 10,500 g at 4°C for 15 mins. Cell pellets (stored at -20°C overnight) were resuspended in 50 mL lysis buffer (350mM NaCl, 20 mM Imidazole, 20 mM Tris pH 8.0, 1mM PMSF, 1 mM EDTA and 3 mM βME) and lysed either via sonication on ice for 3 rounds of 6 mins, 50% duty cycle or using an EmulsiFlex-C3 (AVESTIN, Ontario, Canada), followed by centrifugation at ca. 40,000 g for 1 hour to remove cellular debris. The supernatant was loaded onto a Ni-NTA column equilibrated using 350 mM NaCl, 20 mM Imidazole, 20 mM Tris pH 8.0, 3 mM βME, washed with the same buffer and the SUMO-tagged protein was eluted using 350 mM NaCl, 250 mM Imidazole, 20 mM Tris pH 8.0, 3 mM βME. Protein-containing fractions were pooled and cleaved overnight using SUMO protease (added to final concentration ca. 1 µM), followed by dialysis against 150 mM NaCl, 20 mM Tris pH 8.0, and 1mM DTT and loaded again onto a Ni-NTA column. The cleaved CTDs were collected in the flowthrough and further purified over a 5 mL HiTrap^TM^ SP HP column on an AKTA Pure Protein Purification System (GE) as needed. Purified CTDs were either dialyzed into ddH_2_O, flash frozen and lyophilized or exchanged into NMR Buffer (100 mM NaCl, 10 mM Na_2_HPO_4_ at pH 6.8) by overnight dialysis or using a PD-10 Column (Cytiva, Marlborough, MA) and concentrated as necessary using 3K MWCO Amicon Ultra Centrifugal filters (Millipore Sigma, Darmstadt, Germany) at 10°C. Lyophillized CTDs were resolubilized to ca. 250 uM (by weight) stock concentration in NMR Buffer.

### Large unilamellar vesical (LUV) preparation

18:1 1,2-dioleoyl-sn-glycero-3-phospho-L-serine (DOPS), 18:1 [Δ9-Cis] 1,2-dioleoyl-sn-glycero-3-phosphocholine (DOPC),18:1 [Δ9-Cis] 1,2-dioleoyl-sn-glycero-3-phosphoethanolamine (DOPE), 18:1 1,2-dioleoyl-sn-glycero-3-[{N-(5-amino-1-carboxypentyl)iminodiacetic acid}succinyl] (nickel salt) (DGS-NTA[Ni]), and cholesterol were purchased from Avanti Polar Lipids (Alabaster, AL) dissolved in chloroform and stored at -20 °C. Lipids were mixed to desired ratios (100% DOPS, 1:1 DOPS: DOPC, 100% DOPC, 11:4:5 DOPC:DOPE:DOPS with or without 30% cholesterol, and 10:4:5:1 DOPC:DOPE:DOPS:DGS-Ni-NTA), residual solvent was removed under vacuum for 1-2 hours, and lipids were then stored under N_2_(g) at -20 °C. Lipids were resuspended to 20 mM (total lipid concentration) in 1 mL using either NMR buffer and subjected to 10 freeze/thaw cycles using liquid nitrogen/warm water baths. LUVs were then prepared using a 1 mL Avanti Mini-Extruder (Avanti Polar Lipids) by extruding 21 times using either a 100 nm (for NMR) or 50 nm (for CD) pore size polycarbonate film. Smaller diameter LUVs were used for CD to limit scattering at lower wavelengths that was apparent in data collected for the mGluR3 CTD using NMR samples prepared with 100 nm LUVs (**Fig. S2B**). For vesicle size titrations, LUVs were additionally extruded using 400 nm and 200 nm pore size polycarbonate film. LUVs were stored at 4 °C and used within 5 days.

### Nuclear magnetic resonance (NMR) spectroscopy

CTDs were prepared in NMR Buffer as described above at concentrations ranging from ca. 50-150 µM for 5 mm NMR tubes or 300 µM for 1.7 mm NMR tubes with or without 10 mM LUVs. Relative protein concentrations were corroborated by 1D proton NMR using 4,4-dimethyl-4-silapentane-1-sulfonic acid (DSS) as an internal standard. For lipid titrations LUV concentrations included 10, 5 and 2.5 mM. ^1^H-^15^N HSQC NMR were collected on a Bruker AVANCE 500-MHz spectrometer (Weill Cornell NMR Core) equipped with a Bruker TCI cryoprobe with 1024 complex points in the ^1^H dimension and 146 complex points in the ^15^N dimension using spectral widths of 18 PPM (^1^H) and 24 PPM (^15^N). NMR spectra were acquired at 10 °C to minimize amide protein exchange.

NMR spectra were processed using NMRpipe and analyzed using NMRFAM-sparky 3.115 and NMRbox. Intensity ratios were calculated as the ratio of intensities in spectra of samples with LUVs to those of matched free protein samples with no LUVs. Intensity ratios for cysteine-free CTDs were compared to those of WT CTDs (**Fig. S1C,F**). As no meaningful differences were observed, and as the cysteine residues are located outside the membrane-binding regions of both CTDs, the cysteine-free versions were used in all subsequent experiments. To assess the binding of different CTD regions for LUVs of different compositions and size and for CTD mutants, intensity ratios were averaged over the regions of interest.

Backbone resonance assignments were made using standard triple resonance experiments collected on a Bruker Avance III spectrometer at 800 MHz. Assignments were initially obtained for the WT mGluR2 and mGluR3 CTDs and were transferred by inspection for most mutants, but were augmented using HNCA spectra in select cases (mGluR2 C859A and mGluR3 G848E). Secondary Cα shifts were calculated by subtracting reference random coil Cα shifts^46^ from the observed values.

### Circular dichroism (CD) spectroscopy

CD measurements were performed on an AVIV 410 CD spectropolarimeter. Spectra were obtained from 300-190 nm at 25 °C after a two-minute temperature equilibration with a wavelength step of 1 nm, an averaging time of 5 seconds, 1 scan per sample and a cell path length of 0.02 cm (Starna, Atascadero, CA). Spectra were colletected at a higher temperature compared to NMR spectra to eliminate effects from condensation on the sample cell during data collection. Backgrounds were collected and subtracted from all spectra. Final CTD concentrations ranged from ca. 50-100 µM with protein and peptide stock concentrations estimated by weight, or using A280 with a calculated extinction coefficient of 2980 M^-1^cm^-1^ for mGluR3 CTD, and also assessed using 1D proton NMR with DSS as an internal standard. WT mGluR 2 and mGluR3 peptides encompassing the first 23 residues of each CTD (mGluR2:PQKNVVSHRAPTSRFGSAAPRAS, mGluR3:PQKNVVTHRLHLNRFSVSGTGTT) were purchased from GenScript (Piscataway, NJ) (>= 95% purity). Peptides were dissolved in NMR buffer at 1 mg/mL and mixed 1:1 with CD buffer or LUVs in CD buffer for a final concentration of 0.5 mg/mL. NTS1 peptide encompassing the first 16 residues of its CTD (SANFRQVFLSTLACLC) purchased from from GenScript (>= 95% purity) was dissolved in 100% trifluoroethanol (TFE) at 15 mgs/ml as previously reported^45^ prior to dilution with NMR buffer to 0.4 mg/mL and final 1:1 mixing with buffer or LUVs to a final concentration of 0.2 mg/mL with 0.67% residual TFE. The dearth of aromatic residues in all of the polypeptides used made reliable determination of absolute protein concentrations exceedingly difficult. Therefore, CD data are presented in millidegrees and were not converted to mean residual molar ellipticity.

### Molecular dynamics simulations

All-atom molecular dynamics simulations were performed on an mGluR3 construct containing both TM7 and the CTD (residues 796-879). We used AlphaFold2^75^ and ColabFold^76^ to create models of the mGluR3 TMD and CTD structure because no structures of mGluRs contain a fully resolved CTD. We chose the top ranked of five models (highest average AlphaFold pLDDT score), all of which contained a disordered CTD, and isolated the TM7-CTD residues as our starting structure. CHARMM-GUI^77–83^ was used to build a single system of our starting structure embedded in a 1:1 DOPS-DOPC phospholipid bilayer (139 lipids per leaflet), including 104,130 water molecules and 0.2 M NaCl at 37 °C. This temperature, different from those used for NMR and CD data, was selected to maximize the relevance of the simulations to physiological conditions. The fact that results from these three methods are reasonably self-consistent, despite the use of different temperatures, and are also in agreement with cellular internalization experiments performed at 37 °C, suggests that the fundamentals of mGluR3 CTD-membrane interactions (regions that interact, the driving forces for the interactions, and the lack of secondary structure formation) are not altered over this temperature range to an extent that would alter our results, interpretations or conclusions. Lysine and histidine residues were fully protonated to reflect the decreased pH at the membrane surface^84^.

The initial simulation was equilibrated using the standard CHARMM-GUI based protocol and scripts followed by a short, 6-ns run using OpenMM^85^ and the CHARMM36m^86^ forcefield. Following this equilibration protocol, velocities for each atom in the system were randomized and used to create 6 statistically independent replicates that were then simulated for 1,370 ns for a total of 8.22 µs of simulation time.

Analysis of the trajectories was performed using a combination of VMD^87^ plug-ins and home-made TCL and python scripts. To determine the position of side chains relative to the phosphate plane, we adapted an approach from Marx and Fleming^88^ in which we compared the z-position of a side chain center of mass to the average z-position of the phosphate atoms in the bilayer leaflet closest to the CTD in each frame. Distance from the phosphate plane of the whole CTD or of CTD sub-regions as in Figure S8H was calculated from the center of mass of all side chain and backbone atoms. Side chain-lipid hydrogen bonds were calculated with the Hbonds plug-in in VMD using default settings for bond angle and length. HullRad^89,90^ was used to calculate the radius of gyration of each frame of the simulation. Secondary structure of each residue was calculated for each frame using STRIDE^91^.

### Patch clamp electrophysiology

Whole-cell patch clamp recordings were performed in HEK 293 cells 24 hours post-transfection as previously described^92^. Briefly, cells were transfected using Lipofectamine 2000 (Thermo Fisher Scientific) with the wildtype or mutant mGluRs together with GIRK1-F137S^93^ and soluble tdTomato as a transfection marker in 1:1:0.2 ratio. Recording pipettes of borosilicate and with 3-5 MΩ resistance were filled with intracellular solution (in mM): 140 KCl, 10 HEPES, 5 EGTA, 3 MgCl2, 3 Na_2_ATP, 0.2 Na_2_GTP. Cells were clamped at -60 mV in a high-potassium bath solution containing (in mM): 120 KCl, 25 NaCl, 10 HEPES, 2 CaCl_2_, 1 MgCl_2_. Glutamate evoked potassium currents were recorded with an Axopatch 200B and Digidata 1550B (Molecular Devices) using Clampex (pCLAMP) acquisition software. Peak currents for the different glutamate concentrations were measured using Clampfit (pCLAMP) and normalized to the maximum current observed at saturating 1 mM glutamate.

### Cell imaging and surface labelling assay

To image and quantify the internalization of receptors after glutamate exposure we used N-terminally SNAP-tagged mGluR3 receptors^34^. HEK 293T Cells were transfected with the wildtype or mutant SNAP-mGluR3s. 24 hrs post-transfection cells were prepared for experiments by labeling tem with 1 µM BG-Surface Alexa 546 (non-permeable dye, New England BioLabs) for 30 min at 37°C in a media containing (in mM): 10 HEPES, 135 NaCl, 5.4 KCl, 2 CaCl_2_, 1 MgCl_2_, pH 7.4. After this, cells were washed twice and incubated with glutamate (1 mM) for 30 min. Cells were then imaged under an Olympus IX83 inverted microscope using a 100X 1.49 NA objective and an Orca-Flash 4.0 CMOS camera (Hamamatsu). Representative images were selected over a minimum of 10 images per condition.

For quantifying internalization after glutamate of the wildtype and mutant receptors we used a previously reported surface labelling assay^34^. Briefly, cells expressing different SNAP-tagged mGluR3 constructs were exposed to 1 mM glutamate, or 20 µM of the antagonist LY341495 (Tocris) for normalization, for 1 hr at 37°C. Following drug incubation, cells were labelled with 1 µM BG-Surface Alexa 546 for 20 min at room temperature and imaged immediately using a 60x 1.49 NA objective. Images were analyzed using Image J (Fiji). Mean intensity of thresholded fluorescent areas was calculated using a macro based in the Li algorithm. Background of the tresholded images was subtracted and the resultant values were normalized to the antagonist incubated condition, inverted to show amount of internalization (instead of fluorescence drop) and converted to percentage of internalization. These values were averaged per day. Each point of the bar plots represents a day of experiments. For EGF-stimulated internalization we followed the same approach but applying 100 ng/mL of EGF alone or together with 1 mM Glu. Quantification was performed using the same Li algorithm macro and comparisons were made within each condition (EGF or Glu+EGF) for WT vs Y853A mutant. For the EGFR and pan-Src inhibitors experiment, cells were preincubated for 1 hr with AG1478 (5 µM; Tocris) or Dasatinib (50 µM; Tocris). Afterward, the inhibitors were maintained throughout the whole experiment, which was performed as described above.

---

**Figure S1:**
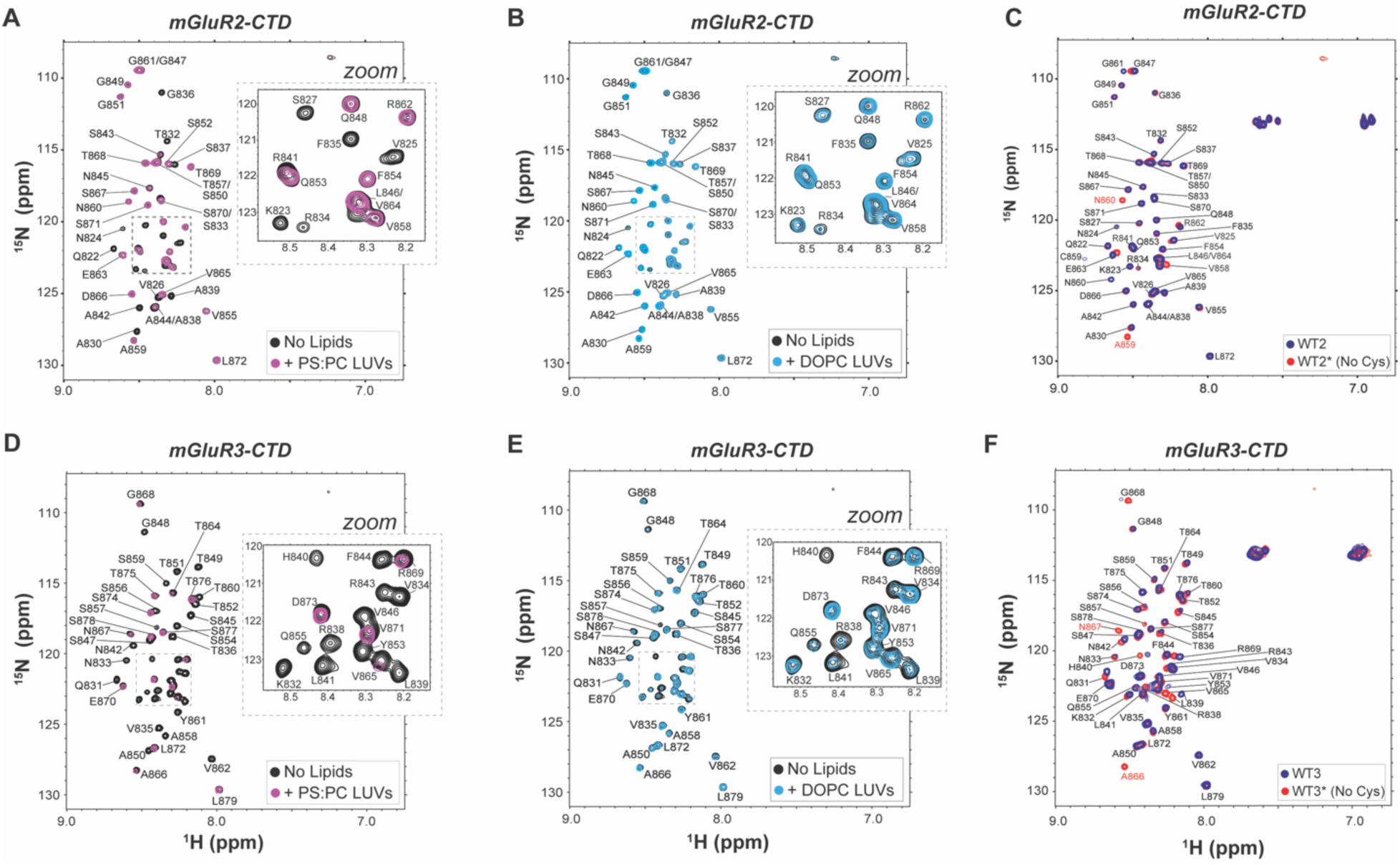
NMR spectra of mGluR2 and mGluR3 with LUVs of different compositions and of WT and cysteine-free variants. ^1^H-^15^N HSQC spectra of mGluR2 **(A,B)** and mGluR3 **(D,E)** CTDs in the presence (purple or light blue) and absence (black) of LUVs comprised of 1:1 DOPS:DOPC **(A,D)** or DOPC **(B,E)** with zoomed insets of crowded regions highlighting loss of signal of specific residues in the presence of LUVs. ^1^H-^15^N HSQC spectra of **(C)** mGluR2 WT (dark blue) and WT* (C859A mutant, red) CTD and **(F)** mGluR3 WT (dark blue) and WT* (C866A mutant, red) CTD. Chemical shift changes are confined to residues in the immediate vicinity of the mutations.

**Figure S2:**
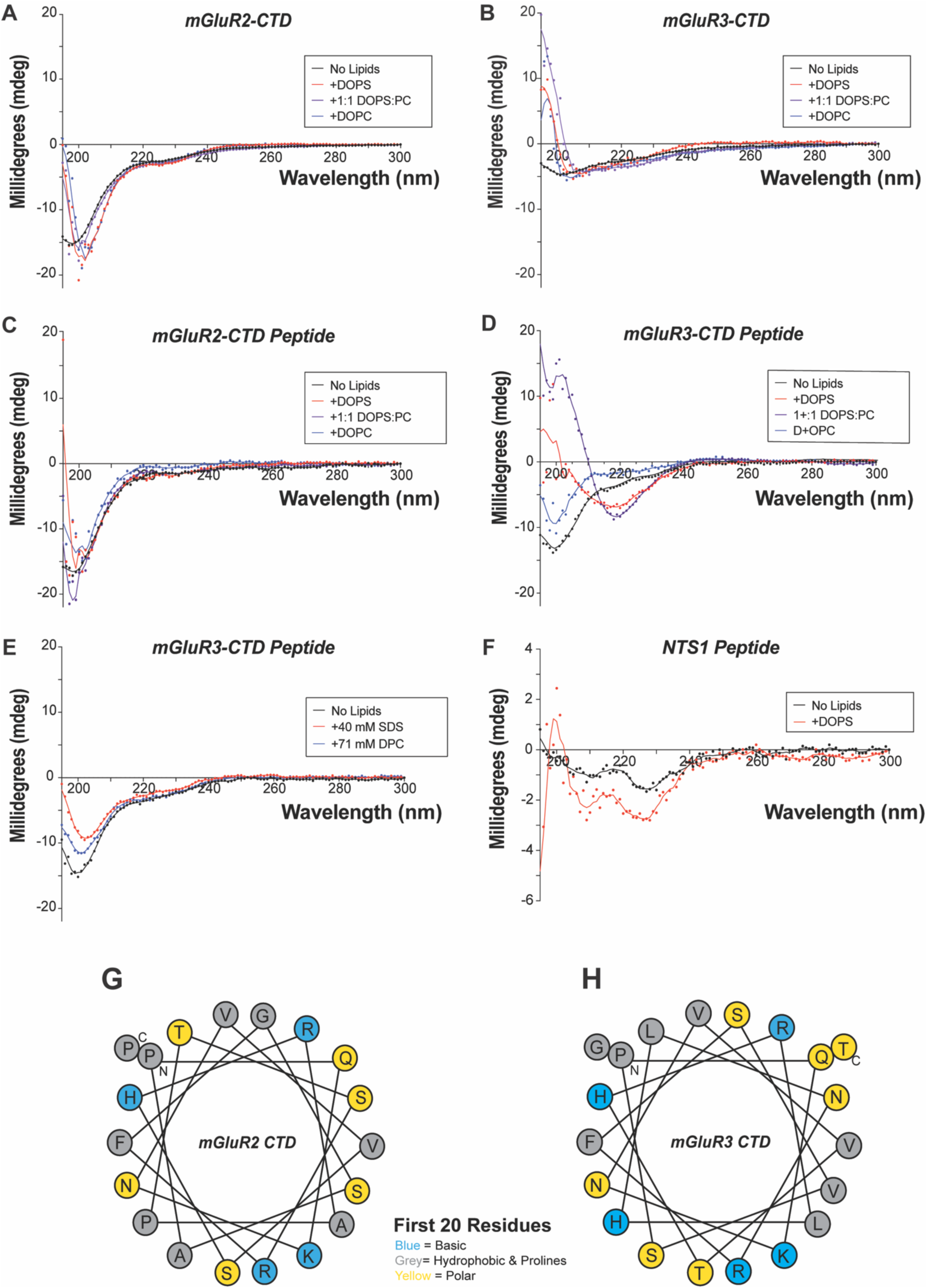
Secondary structure analysis of mGluR CTDs using CD. Circular dichroism spectra of the mGluR2 **(A)** and mGluR3 **(B)** CTD and mGluR2 **(C)** and mGluR3 **(D)** peptides encompassing the NMR determined binding region (first 23 residues) in the absence of LUVs (black) or in the presence of 10 mM, 100 nm diameter **(B)** or 50 nm diameter **(A,C,D)** DOPC (blue), 1:1 DOPS:DOPC (purple) or DOPS (red) LUVs under conditions where NMR indicates no (DOPC) or complete (DOPS or 1:1 DOPS:DOPC) CTD binding. Note that for **(B)** the sudden increase in signal below 205 nm results from increased scattering due to the use of 100 nm diameter LUVs in these samples. For **(D)** the spectra suggest a degree of β-strand formation, possibly due to intermolecular interactions caused by the limited solubility of this peptide **(E)** CD spectra of mGluR3 peptides encompassing the NMR determined binding region (first 23 residues) in the absence (black) or presence of 40 mM SDS (red) or 71 mM DPC (blue). **(F)** CD spectra of NTSR1 peptide free in solution (black) and in the presence of 10 mM 50 nm diameter DOPS LUVs (red). **(G,H)** Helical wheel representations of the first 20 residues of the mGluR2 **(G)** and mGluR3 **(H)** CTD illustrating the lack of clear amphipathic character (basic residue in blue, polar residues in yellow, apolar residues in grey).

**Figure S3:**
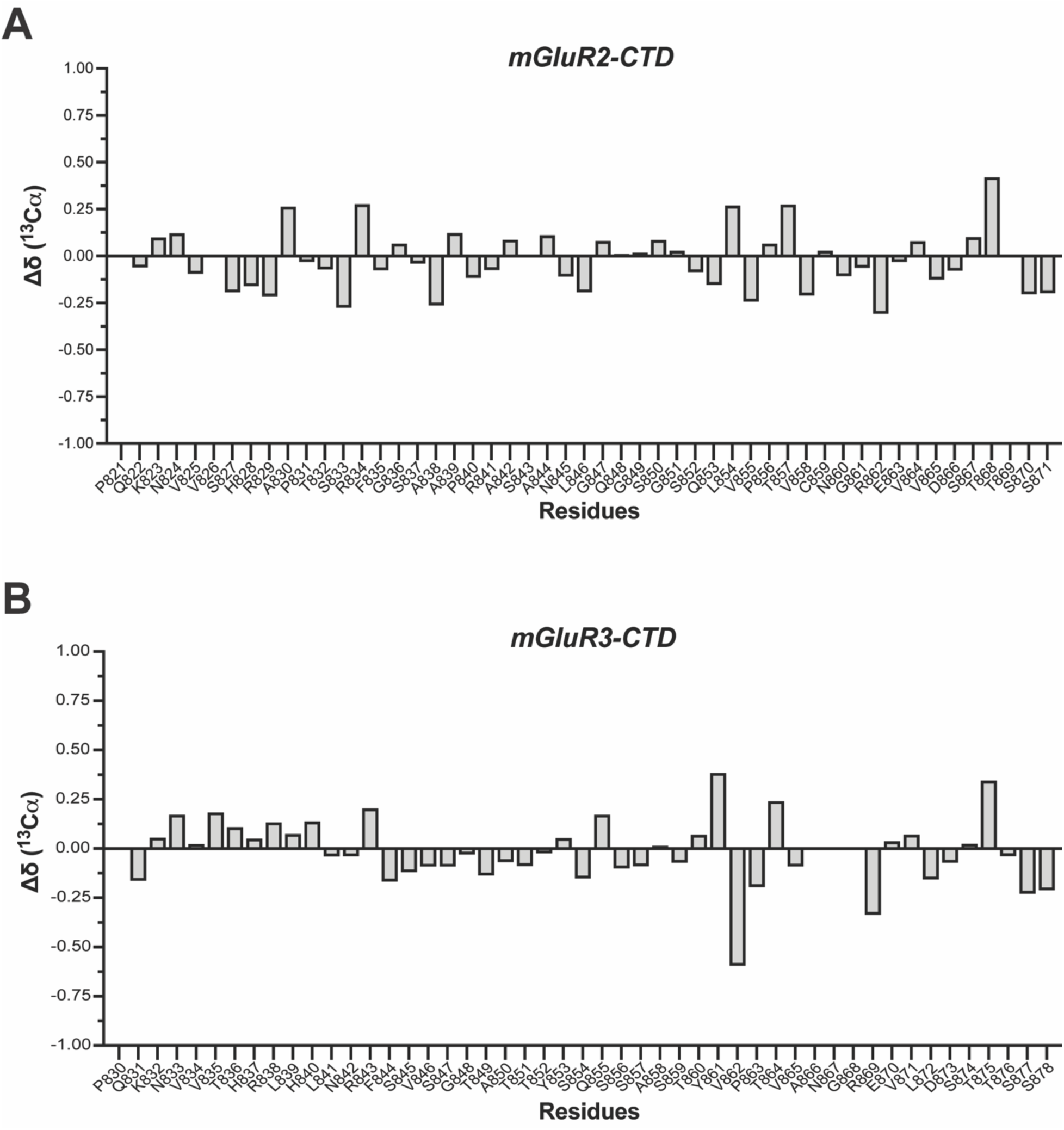
Secondary structure analysis of mGluR CTDs using NMR. Secondary C_α_ chemical shifts for the mGluR2 **(A)** and mGluR3 **(B)** CTDs free in solution. Positive/negative deviations greater than 1.0 PPM indicate a propensity for helical/μ-strand secondary structure, respectively.

**Figure S4:**
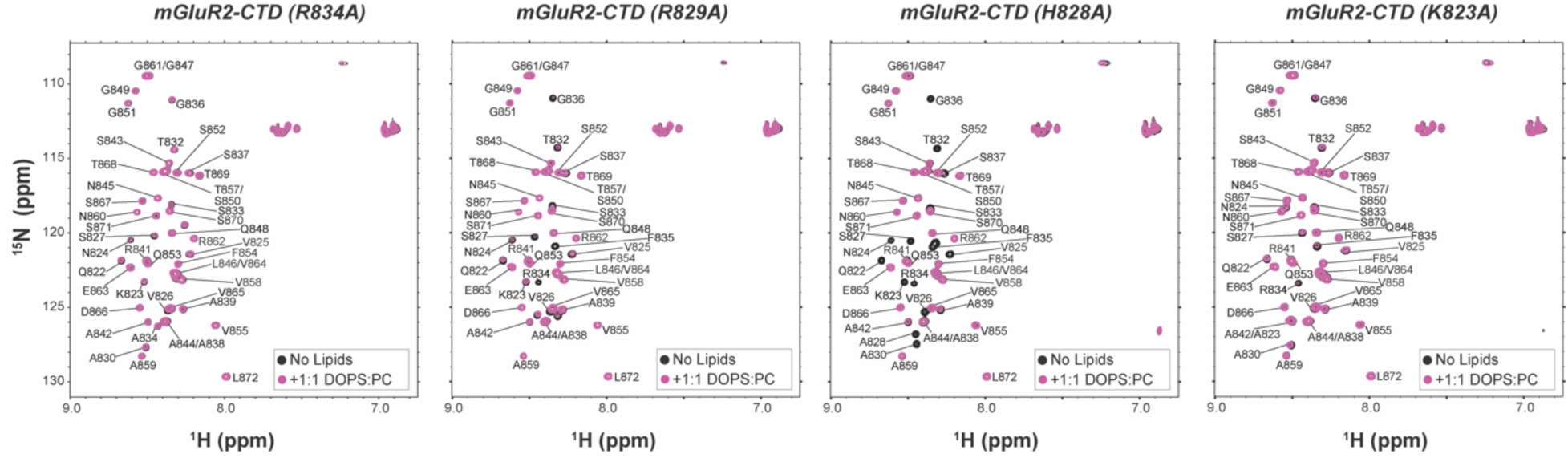
Mutating positively charged residues impacts CTD-membrane interactions. ^1^H-^15^N HSQC spectra of mGluR2 CTD variants in the presence (purple) and absence (black) of 100 nm LUVs comprised of 1:1 DOPS:DOPC.

**Figure S5:**
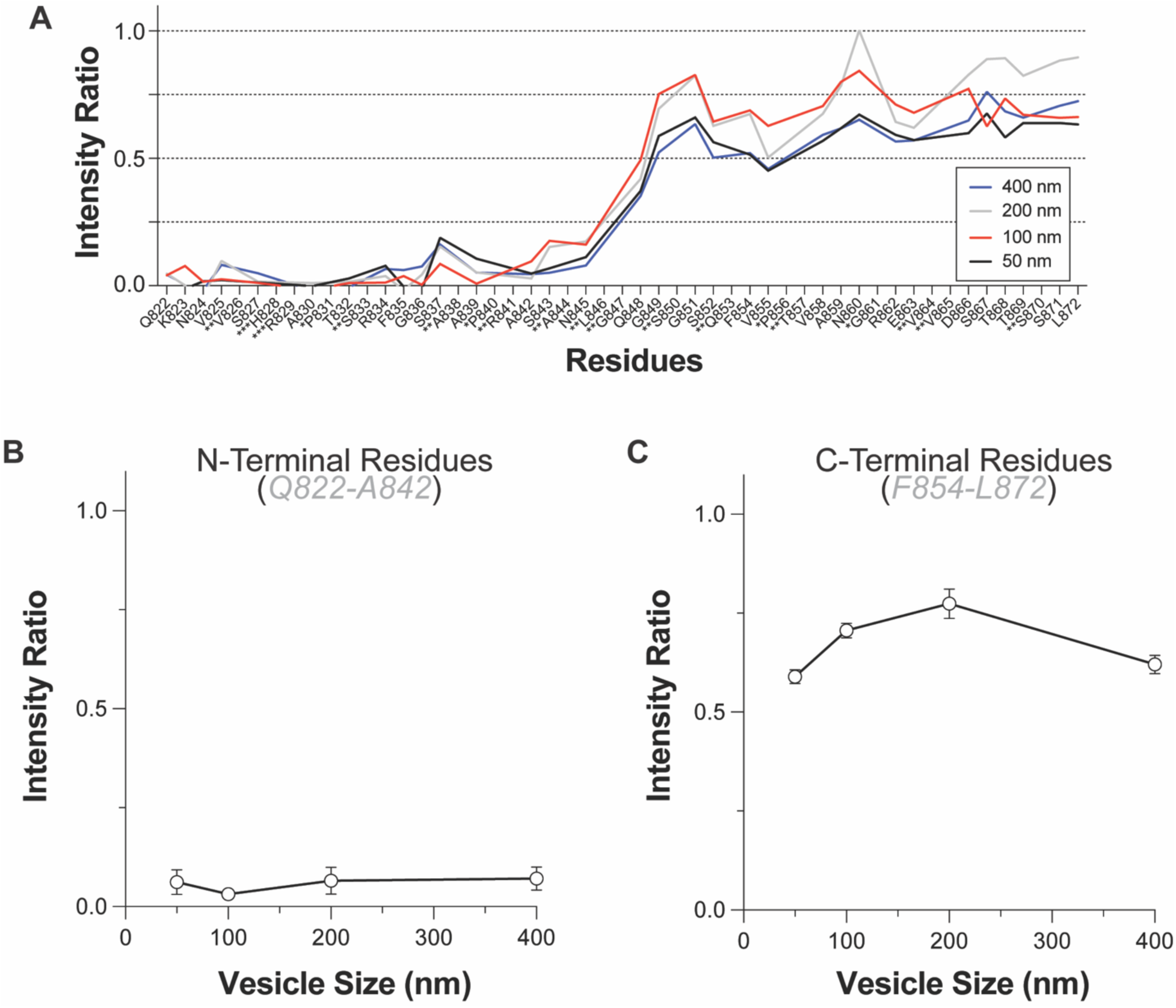
mGluR CTDs LUVs binding as a function of vesicle size. **(A)** NMR intensity ratio plots for the mGluR2-CTD in the presence of DOPS LUVs of varying diameter. **(B)** Averaged intensity ratios over the first ∼20 residues (Q822-A842) (±s.e.m.). **(C)** Averaged intensity ratios over the last ∼20 residues (Q853-L872) (±s.e.m.).

**Figure S6:**
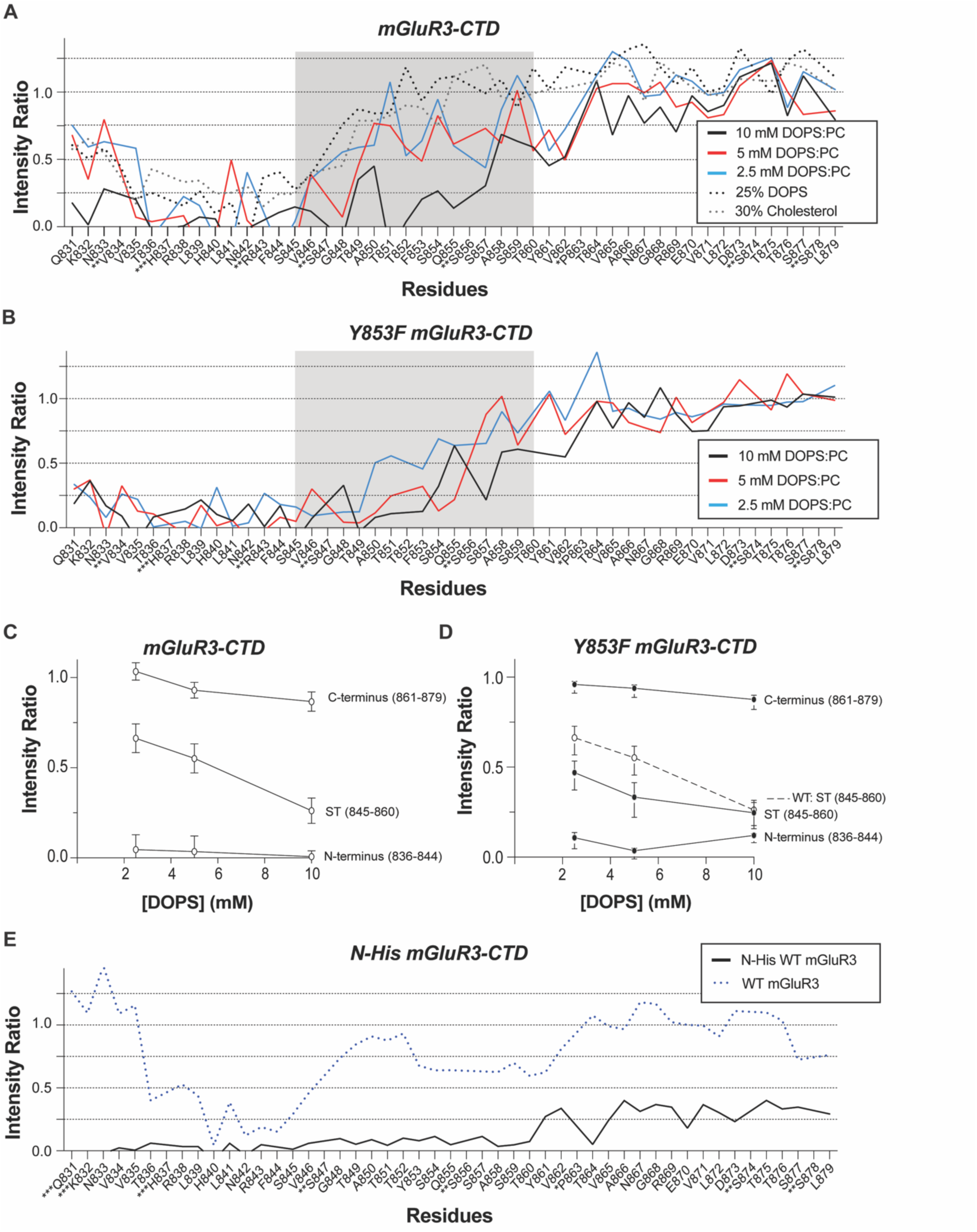
Lipid titrations identify weaker and stronger interaction regions of WT and Y853F mGluR3 CTD. **(A)** NMR intensity ratios for the WT mGluR3-CTD in the presence of varying concentrations of 1:1 DOPS:DOPC LUVs and of 10 mM 11:4:5 DOPC:DOPE:DOPS with or without 30% cholesterol. **(B)** NMR intensity ratios for the Y853F mGluR3-CTD in the presence of varying concentrations of 1:1 DOPS:DOPC LUVs. **(C)** Average intensity ratios over different regions of the WT mGluR3 CTD highlight the greater sensitivity of the S/T-rich region to lipid concentration (±s.e.m.). **(D)** Average intensity ratios over different regions of the Y853F mGluR3 CTD highlight the increased binding of the S/T-rich region at lower lipid concentration compared to WT (±s.e.m.). **(E)** NMR intensity ratios for the membrane-tethered 6x N-His WT mGluR3-CTD (black) compared to untethered WT mGluR3-CTD (dotted blue) in the presence of 10 mM 10:4:5:1 DOPC:DOPE:DOPS:DGS-Ni-NTA LUVs.

**Figure S7:**
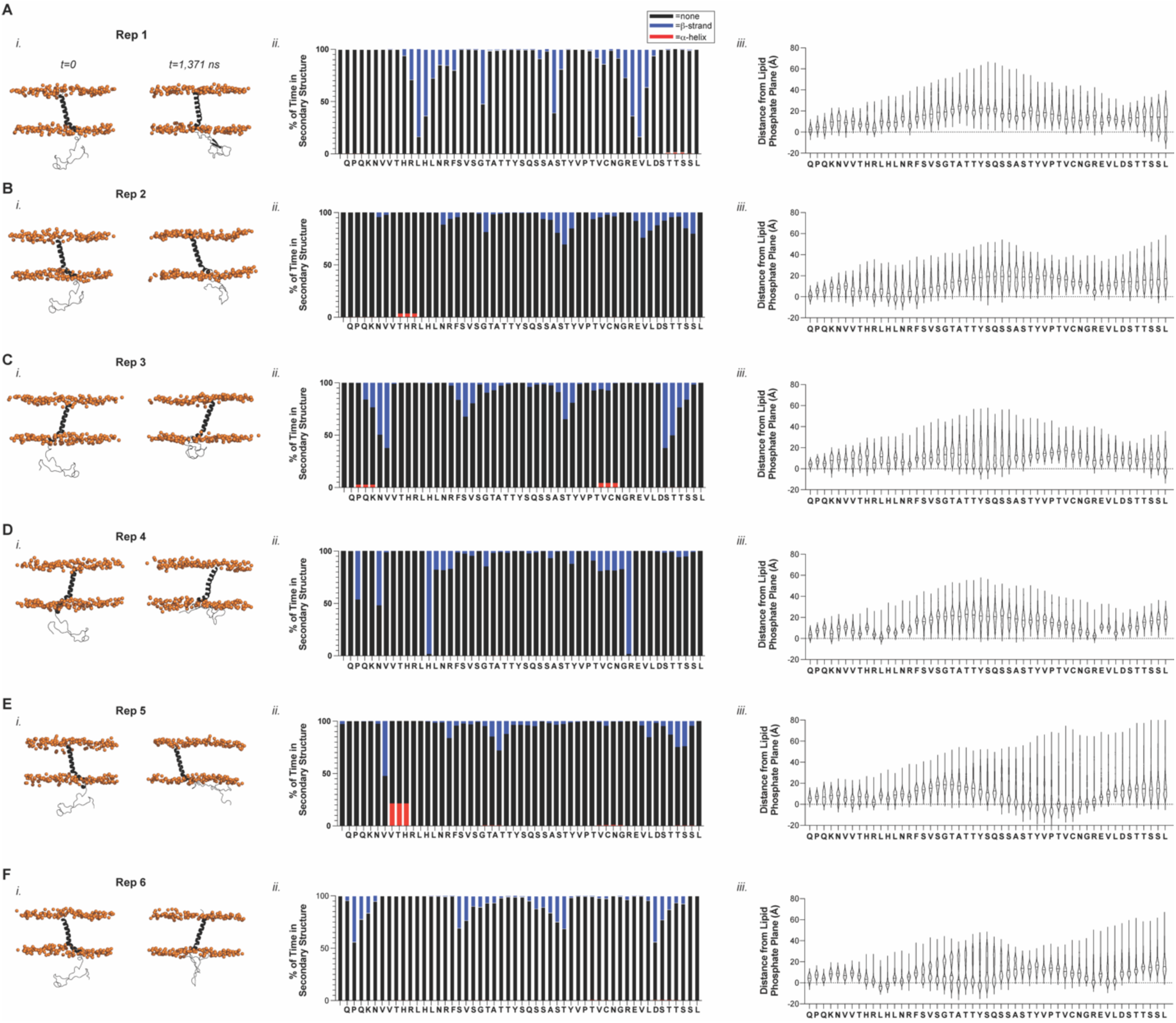
MD simulations capture mGluR3 CTD-membrane interactions and confirm absence of regular secondary structure. (**A-F)** (i, left) snapshots of starting and ending frames of each MD simulation replica of the mGluR3 TM7-CTD construct with the protein shown in black cartoon representation and lipid phosphates as orange spheres; (ii, middle) Percent of trajectory frames for each CTD residue residing in disordered (black), helical (red), or beta-sheet (blue) secondary structure during each replica trajectory.; (iii, right) Violin plots of average residue distance from the phosphate plane in each replica.

**Figure S8:**
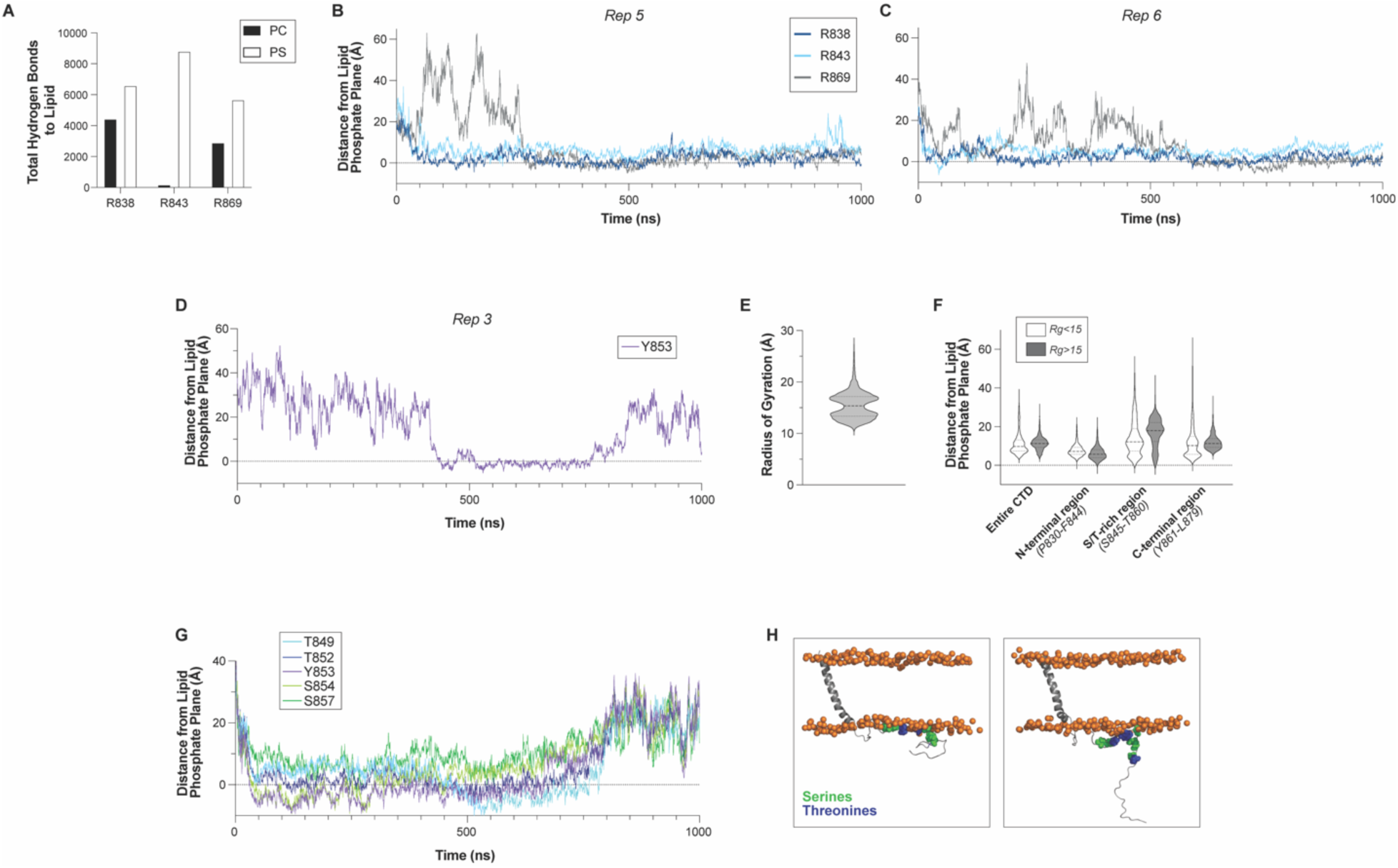
Comparison and analysis of MD simulation replicas. **(A)** Total number of hydrogen bonds summed over all MD replicas between key arginine side chains and DOPC or DOPS lipid headgroups, highlighting the preference for interactions with negatively charged PS headgroups**. (B, C)** Position of arginine side chains relative to the phosphate plane of the membrane for the first 1,000 ns of MD replica 5 **(B)** and 6 **(C)**. **(D)** Position of tyrosine 853 side chain relative to the phosphate plane of the membrane for the first 1,000 ns of MD replica 2. **(E)** Distance from phosphate plane of side chains in the S/T-rich region as a function of time for Y853 and nearby S/T residues in MD replica 6 (first 1,000 ns). **(F)** Snapshots of S/T residues in the S/T-rich region relative to the membrane highlighting the dynamics of the region during our simulations. Side chains are shown as spheres (Ser in green and Thr in blue) and lipid phosphates are shown as orange spheres. Protein backbone is in gray cartoon. **(G)** Violin plot of radii of gyration for all MD replicas combined showing pinch point at 15 Å. **(H)** Violin plots showing distributon of distances to the membrane for the entire CTD and specific CTD subregions for frames with high (>15 Å) and low (<15 Å) Rg.

**Figure S9:**
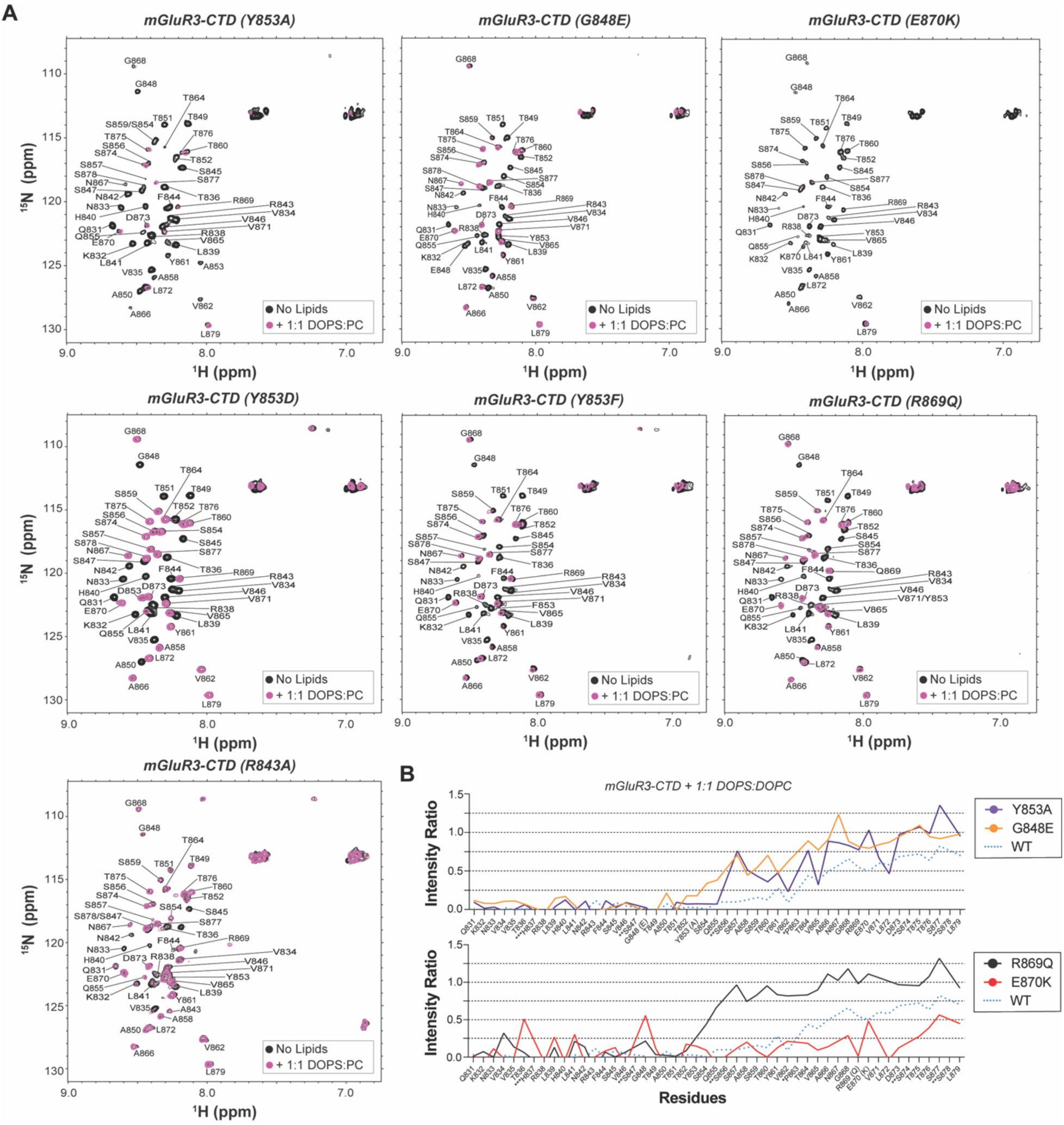
De novo and cancer-associated mutations modulate membrane binding of the mGlur3 S/T-rich region. **(A)** ^1^H-^15^N HSQC spectra of mGluR3 CTD variants in the presence (purple) and absence (black) of 10 mM 1:1 DOPS:DOPC LUVs. **(B)** NMR intensity ratios for WT (dotted blue line), Y853A (purple) and G848E (orange) [upper panel] and WT (dotted blue line), R869Q (black) and E870K (red) [lower panel] mGluR3-CTD in the presence of 10 mM 1:1 DOPS:DOPC LUVs.

**Figure S10:**
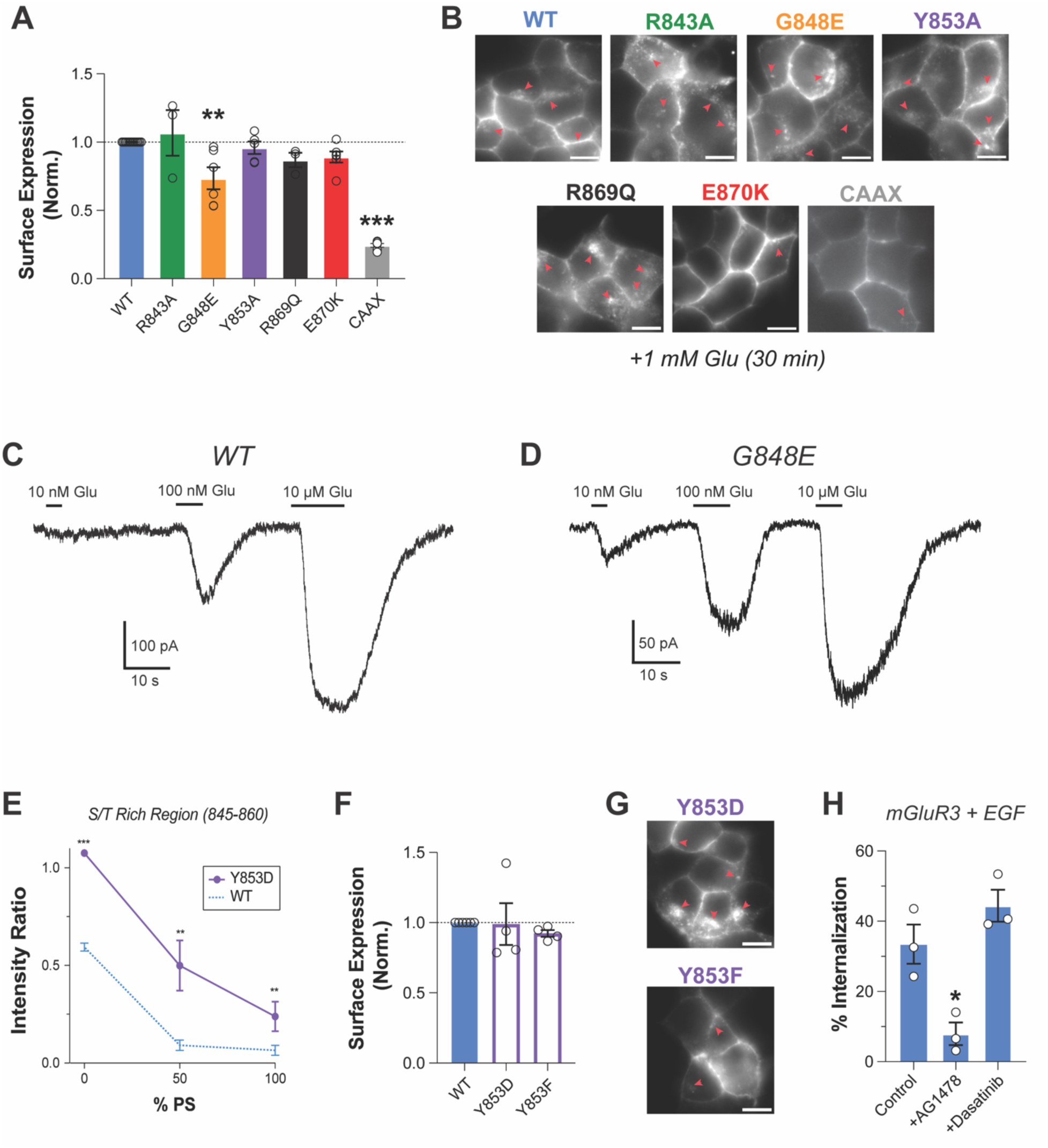
Further analysis of CTD mutants and effects on mGluR3 function. **(A)** Surface expression quantification for R843A, G848E, Y853A, R869Q, E870K and -CAAX mGluR3 (One-way ANOVA with multiple comparisons; * p<0.05, *** p<0.001). **(B)** Representative images of HEK 293T cells expressing SNAP-tagged mGluR3 variants incubated for 30 min with 1 mM Glu (red arrows represent internalized receptors; scale bar: 5 µm). **(C-D)** Representative traces of evoked GIRK potassium currents after activation by different doses of glutamate (10 nM, 100 nM and 10 µM) for WT **(C)** and G848E **(D)** mGluR3**. (E)** Comparison of the averaged integrated NMR intensity ratios of WT mGluR3-CTD (dotted blue) with Y853D (phosphomimetic) mGluR3-CTD taken over the S/T-rich region (S845-T860) as a function of LUV lipid composition (±s.e.m.; Wilcoxon test; *p<0.05, ***p<0.001, n.s. p≥0.05). **(F)** Surface expression quantification for Y853D and Y853F mGluR3 (One-way ANOVA with multiple comparisons). **(G)** Representative images of HEK 293T cells expressing SNAP-tagged mGluR3 Y853D (upper panel) and Y853F (lower panel) incubated for 30 min with 1 mM Glu (red arrows represent internalized receptors; scale bar: 5 µm). **(H)** Inhibition of EGF-induced internalization with the EGFR tyrosine kinase inhibitor AG1478 (5 µM) or the pan-Src family inhibitor Dasatinib (50 µM), (One-way ANOVA with multiple comparisons; *p<0.05).

## References

1 Wacker, D., Stevens, R. C. & Roth, B. L. How Ligands Illuminate GPCR Molecular Pharmacology. Cell 170, 414–427, doi:10.1016/j.cell.2017.07.009 (2017).

2 Sriram, K. & Insel, P. A. G Protein-Coupled Receptors as Targets for Approved Drugs: How Many Targets and How Many Drugs? Mol Pharmacol 93, 251–258, doi:10.1124/mol.117.111062 (2018).

3 Guillien, M. et al. Structural Insights into the Intrinsically Disordered GPCR C-Terminal Region, Major Actor in Arrestin-GPCR Interaction. Biomolecules 12, doi:10.3390/biom12050617 (2022).

4. Guillien, M. et al. Phosphorylation motif dictates GPCR C-terminal domain conformation and arrestin interaction. bioRxiv, 2023.2002.2023.529712, doi:10.1101/2023.02.23.529712 (2023).

5 Heng, J. et al. Function and dynamics of the intrinsically disordered carboxyl terminus of beta2 adrenergic receptor. Nat Commun 14, 2005, doi:10.1038/s41467-023-37233-1 (2023).

6 Shiraishi, Y. et al. Phosphorylation-induced conformation of beta(2)-adrenoceptor related to arrestin recruitment revealed by NMR. Nat Commun 9, 194, doi:10.1038/s41467-017-02632-8 (2018).

7 Maeda, S., Qu, Q., Robertson, M. J., Skiniotis, G. & Kobilka, B. K. Structures of the M1 and M2 muscarinic acetylcholine receptor/G-protein complexes. Science 364, 552–557, doi:10.1126/science.aaw5188 (2019).

8 Tsai, C. J. et al. Cryo-EM structure of the rhodopsin-Galphai-betagamma complex reveals binding of the rhodopsin C-terminal tail to the gbeta subunit. Elife 8, doi:10.7554/eLife.46041 (2019).

9 Seven, A. B. et al. G-protein activation by a metabotropic glutamate receptor. Nature 595, 450–454, doi:10.1038/s41586-021-03680-3 (2021).

10 He, F. et al. Allosteric modulation and G-protein selectivity of the Ca(2+)-sensing receptor. Nature 626, 1141–1148, doi:10.1038/s41586-024-07055-2 (2024).

11 Kisselev, O. G., McDowell, J. H. & Hargrave, P. A. The arrestin-bound conformation and dynamics of the phosphorylated carboxy-terminal region of rhodopsin. FEBS Lett 564, 307–311, doi:10.1016/S0014-5793(04)00226-1 (2004).

12 Shukla, A. K. et al. Structure of active beta-arrestin-1 bound to a G-protein-coupled receptor phosphopeptide. Nature 497, 137–141, doi:10.1038/nature12120 (2013).

13 Min, K. et al. Crystal Structure of beta-Arrestin 2 in Complex with CXCR7 Phosphopeptide. Structure 28, 1014–1023 e1014, doi:10.1016/j.str.2020.06.002 (2020).

14 Zhou, X. E. et al. Identification of Phosphorylation Codes for Arrestin Recruitment by G Protein-Coupled Receptors. Cell 170, 457–469 e413, doi:10.1016/j.cell.2017.07.002 (2017).

15 Yin, W. et al. A complex structure of arrestin-2 bound to a G protein-coupled receptor. Cell Res 29, 971–983, doi:10.1038/s41422-019-0256-2 (2019).

16 Staus, D. P. et al. Structure of the M2 muscarinic receptor-beta-arrestin complex in a lipid nanodisc. Nature 579, 297–302, doi:10.1038/s41586-020-1954-0 (2020).

17 Huang, W. et al. Structure of the neurotensin receptor 1 in complex with beta-arrestin 1. Nature 579, 303–308, doi:10.1038/s41586-020-1953-1 (2020).

18 Ahn, K. H. et al. Structural analysis of the human cannabinoid receptor one carboxyl-terminus identifies two amphipathic helices. Biopolymers 91, 565–573, doi:10.1002/bip.21179 (2009).

19 Patwardhan, A., Cheng, N. & Trejo, J. Post-Translational Modifications of G Protein-Coupled Receptors Control Cellular Signaling Dynamics in Space and Time. Pharmacol Rev 73, 120–151, doi:10.1124/pharmrev.120.000082 (2021).

20 Song, W., Yen, H. Y., Robinson, C. V. & Sansom, M. S. P. State-dependent Lipid Interactions with the A2a Receptor Revealed by MD Simulations Using In Vivo-Mimetic Membranes. Structure 27, 392–403 e393, doi:10.1016/j.str.2018.10.024 (2019).

21 Dawaliby, R. et al. Allosteric regulation of G protein-coupled receptor activity by phospholipids. Nat Chem Biol 12, 35–39, doi:10.1038/nchembio.1960 (2016).

22 Yen, H. Y. et al. PtdIns(4,5)P2 stabilizes active states of GPCRs and enhances selectivity of G-protein coupling. Nature 559, 423–427, doi:10.1038/s41586-018-0325-6 (2018).

23 Janetzko, J. et al. Membrane phosphoinositides regulate GPCR-beta-arrestin complex assembly and dynamics. Cell 185, 4560–4573 e4519, doi:10.1016/j.cell.2022.10.018 (2022).

24. Grimes, J. et al. Single-molecule analysis of receptor-β-arrestin interactions in living cells. *bioRxiv*, 2022.2011.2015.516577, doi:10.1101/2022.11.15.516577 (2022).

25. Chen, Q. et al. ACKR3-arrestin2/3 complexes reveal molecular consequences of GRK-dependent barcoding. *bioRxiv*, doi:10.1101/2023.07.18.549504 (2023).

26 Ellaithy, A., Gonzalez-Maeso, J., Logothetis, D. A. & Levitz, J. Structural and Biophysical Mechanisms of Class C G Protein-Coupled Receptor Function. Trends Biochem Sci 45, 1049–1064, doi:10.1016/j.tibs.2020.07.008 (2020).

27 Dore, A. S. et al. Structure of class C GPCR metabotropic glutamate receptor 5 transmembrane domain. Nature 511, 557–562, doi:10.1038/nature13396 (2014).

28 Wu, H. et al. Structure of a class C GPCR metabotropic glutamate receptor 1 bound to an allosteric modulator. Science 344, 58–64, doi:10.1126/science.1249489 (2014).

29 Du, J. et al. Structures of human mGlu2 and mGlu7 homo-and heterodimers. Nature 594, 589–593, doi:10.1038/s41586-021-03641-w (2021).

30 Enz, R. Structure of metabotropic glutamate receptor C-terminal domains in contact with interacting proteins. Front Mol Neurosci 5, 52, doi:10.3389/fnmol.2012.00052 (2012).

31 Reiner, A. & Levitz, J. Glutamatergic Signaling in the Central Nervous System: Ionotropic and Metabotropic Receptors in Concert. Neuron 98, 1080–1098, doi:10.1016/j.neuron.2018.05.018 (2018).

32 Pin, J. P. & Bettler, B. Organization and functions of mGlu and GABA(B) receptor complexes. Nature 540, 60–68, doi:10.1038/nature20566 (2016).

33 Suh, Y. H., Chang, K. & Roche, K. W. Metabotropic glutamate receptor trafficking. Mol Cell Neurosci 91, 10–24, doi:10.1016/j.mcn.2018.03.014 (2018).

34 Abreu, N., Acosta-Ruiz, A., Xiang, G. & Levitz, J. Mechanisms of differential desensitization of metabotropic glutamate receptors. Cell Rep 35, 109050, doi:10.1016/j.celrep.2021.109050 (2021).

35 Lin, S. et al. Structures of G(i)-bound metabotropic glutamate receptors mGlu2 and mGlu4. Nature 594, 583–588, doi:10.1038/s41586-021-03495-2 (2021).

36 Koehl, A. et al. Structural insights into the activation of metabotropic glutamate receptors. Nature 566, 79–84, doi:10.1038/s41586-019-0881-4 (2019).

37 Nasrallah, C. et al. Agonists and allosteric modulators promote signaling from different metabotropic glutamate receptor 5 conformations. Cell Rep 36, 109648, doi:10.1016/j.celrep.2021.109648 (2021).

38. Strauss, A. et al. Structural basis of allosteric modulation of metabotropic glutamate receptor activation and desensitization. *bioRxiv*, 2023.2008.2013.552748, doi:10.1101/2023.08.13.552748 (2023).

39 Das, T. & Eliezer, D. Membrane interactions of intrinsically disordered proteins: The example of alpha-synuclein. Biochim Biophys Acta Proteins Proteom 1867, 879–889, doi:10.1016/j.bbapap.2019.05.001 (2019).

40 Das, T. & Eliezer, D. Probing Structural Changes in Alpha-Synuclein by Nuclear Magnetic Resonance Spectroscopy. Methods Mol Biol 1948, 157–181, doi:10.1007/978-1-4939-9124-2_13 (2019).

41 Snead, D. & Eliezer, D. Spectroscopic Characterization of Structure-Function Relationships in the Intrinsically Disordered Protein Complexin. Methods Enzymol 611, 227–286, doi:10.1016/bs.mie.2018.08.005 (2018).

42 Parker, W. & Song, P. S. Protein structures in SDS micelle-protein complexes. Biophys J 61, 1435–1439, doi:10.1016/S0006-3495(92)81949-5 (1992).

43 Bussell, R., Jr. & Eliezer, D. A structural and functional role for 11-mer repeats in alpha-synuclein and other exchangeable lipid binding proteins. J Mol Biol 329, 763–778 (2003).

44 Snead, D. et al. Unique Structural Features of Membrane-Bound C-Terminal Domain Motifs Modulate Complexin Inhibitory Function. Front Mol Neurosci 10, 154, doi:10.3389/fnmol.2017.00154 (2017).

45 Dijkman, P. M. et al. Conformational dynamics of a G protein-coupled receptor helix 8 in lipid membranes. Sci Adv 6, eaav8207, doi:10.1126/sciadv.aav8207 (2020).

46 Nielsen, J. T. & Mulder, F. A. A. CheSPI: chemical shift secondary structure population inference. J Biomol NMR 75, 273–291, doi:10.1007/s10858-021-00374-w (2021).

47 Snead, D., Wragg, R. T., Dittman, J. S. & Eliezer, D. Membrane curvature sensing by the C-terminal domain of complexin. Nat Commun 5, 4955, doi:10.1038/ncomms5955 (2014).

48 Antonny, B. Mechanisms of membrane curvature sensing. Annu Rev Biochem 80, 101–123, doi:10.1146/annurev-biochem-052809-155121 (2011).

49 Ingolfsson, H. I. et al. Lipid organization of the plasma membrane. J Am Chem Soc 136, 14554–14559, doi:10.1021/ja507832e (2014).

50 Klahn, M. & Zacharias, M. Transformations in plasma membranes of cancerous cells and resulting consequences for cation insertion studied with molecular dynamics. Phys Chem Chem Phys 15, 14427–14441, doi:10.1039/c3cp52085d (2013).

51 McDonald, S. K. & Fleming, K. G. Aromatic Side Chain Water-to-Lipid Transfer Free Energies Show a Depth Dependence across the Membrane Normal. J Am Chem Soc 138, 7946–7950, doi:10.1021/jacs.6b03460 (2016).

52 Moon, C. P. & Fleming, K. G. Side-chain hydrophobicity scale derived from transmembrane protein folding into lipid bilayers. Proc Natl Acad Sci U S A 108, 10174–10177, doi:10.1073/pnas.1103979108 (2011).

53 Killian, J. A. & von Heijne, G. How proteins adapt to a membrane-water interface. Trends Biochem Sci 25, 429–434, doi:10.1016/s0968-0004(00)01626-1 (2000).

54 Prickett, T. D. et al. Exon capture analysis of G protein-coupled receptors identifies activating mutations in GRM3 in melanoma. Nat Genet 43, 1119–1126, doi:10.1038/ng.950 (2011).

55 Sondka, Z. et al. COSMIC: a curated database of somatic variants and clinical data for cancer. Nucleic Acids Res 52, D1210–D1217, doi:10.1093/nar/gkad986 (2024).

56 Asher, W. B. et al. GPCR-mediated beta-arrestin activation deconvoluted with single-molecule precision. Cell 185, 1661–1675 e1616, doi:10.1016/j.cell.2022.03.042 (2022).

57 Xu, C. et al. Regulation of T cell receptor activation by dynamic membrane binding of the CD3epsilon cytoplasmic tyrosine-based motif. Cell 135, 702–713, doi:10.1016/j.cell.2008.09.044 (2008).

58 Wang, Y. et al. Regulation of EGFR nanocluster formation by ionic protein-lipid interaction. Cell Res 24, 959–976, doi:10.1038/cr.2014.89 (2014).

59 Cornish, J., Chamberlain, S. G., Owen, D. & Mott, H. R. Intrinsically disordered proteins and membranes: a marriage of convenience for cell signalling? Biochem Soc Trans 48, 2669–2689, doi:10.1042/BST20200467 (2020).

60 Lee, J. et al. Distinct beta-arrestin coupling and intracellular trafficking of metabotropic glutamate receptor homo- and heterodimers. Sci Adv 9, eadi8076, doi:10.1126/sciadv.adi8076 (2023).

61 Fang, W. et al. Structural basis of the activation of metabotropic glutamate receptor 3. Cell Res 32, 695–698, doi:10.1038/s41422-022-00623-z (2022).

62 Morton, L. A. et al. MARCKS-ED peptide as a curvature and lipid sensor. ACS Chem Biol 8, 218–225, doi:10.1021/cb300429e (2013).

63 Wang, J. et al. Lateral sequestration of phosphatidylinositol 4,5-bisphosphate by the basic effector domain of myristoylated alanine-rich C kinase substrate is due to nonspecific electrostatic interactions. J Biol Chem 277, 34401–34412, doi:10.1074/jbc.M203954200 (2002).

64 Wragg, R. T. et al. Evolutionary Divergence of the C-terminal Domain of Complexin Accounts for Functional Disparities between Vertebrate and Invertebrate Complexins. Front Mol Neurosci 10, 146, doi:10.3389/fnmol.2017.00146 (2017).

65 Hicks, A., Escobar, C. A., Cross, T. A. & Zhou, H. X. Fuzzy Association of an Intrinsically Disordered Protein with Acidic Membranes. JACS Au 1, 66–78, doi:10.1021/jacsau.0c00039 (2021).

66 Dikiy, I. et al. Semisynthetic and in Vitro Phosphorylation of Alpha-Synuclein at Y39 Promotes Functional Partly Helical Membrane-Bound States Resembling Those Induced by PD Mutations. ACS Chem Biol 11, 2428–2437, doi:10.1021/acschembio.6b00539 (2016).

67 Acosta, D. M., Mancinelli, C., Bracken, C. & Eliezer, D. Post-translational modifications within tau paired helical filament nucleating motifs perturb microtubule interactions and oligomer formation. J Biol Chem 298, 101442, doi:10.1016/j.jbc.2021.101442 (2022).

68 Kilpatrick, L. E. & Hill, S. J. Transactivation of G protein-coupled receptors (GPCRs) and receptor tyrosine kinases (RTKs): Recent insights using luminescence and fluorescence technologies. Curr Opin Endocr Metab Res 16, 102–112, doi:10.1016/j.coemr.2020.10.003 (2021).

69 Chen, Y., Long, H., Wu, Z., Jiang, X. & Ma, L. EGF transregulates opioid receptors through EGFR-mediated GRK2 phosphorylation and activation. Mol Biol Cell 19, 2973–2983, doi:10.1091/mbc.e07-10-1058 (2008).

70 Tobin, A. B. G-protein-coupled receptor phosphorylation: where, when and by whom. Br J Pharmacol 153 **Suppl 1**, S167–176, doi:10.1038/sj.bjp.0707662 (2008).

71 Nobles, K. N. et al. Distinct phosphorylation sites on the beta(2)-adrenergic receptor establish a barcode that encodes differential functions of beta-arrestin. Sci Signal 4, ra51, doi:10.1126/scisignal.2001707 (2011).

72 Gao, Y. et al. Asymmetric activation of the calcium-sensing receptor homodimer. Nature 595, 455–459, doi:10.1038/s41586-021-03691-0 (2021).

73 Pluhackova, K., Wilhelm, F. M. & Muller, D. J. Lipids and Phosphorylation Conjointly Modulate Complex Formation of beta(2)-Adrenergic Receptor and beta-arrestin2. Front Cell Dev Biol 9, 807913, doi:10.3389/fcell.2021.807913 (2021).

74 Wang, X. et al. Structural insights into dimerization and activation of the mGlu2-mGlu3 and mGlu2-mGlu4 heterodimers. Cell Res, doi:10.1038/s41422-023-00830-2 (2023).

75 Jumper, J. et al. Highly accurate protein structure prediction with AlphaFold. Nature 596, 583–589, doi:10.1038/s41586-021-03819-2 (2021).

76 Mirdita, M. et al. ColabFold: making protein folding accessible to all. Nat Methods 19, 679–682, doi:10.1038/s41592-022-01488-1 (2022).

77 Jo, S., Kim, T., Iyer, V. G. & Im, W. CHARMM-GUI: a web-based graphical user interface for CHARMM. J Comput Chem 29, 1859–1865, doi:10.1002/jcc.20945 (2008).

78 Brooks, B. R. et al. CHARMM: the biomolecular simulation program. J Comput Chem 30, 1545–1614, doi:10.1002/jcc.21287 (2009).

79 Lee, J. et al. CHARMM-GUI Input Generator for NAMD, GROMACS, AMBER, OpenMM, and CHARMM/OpenMM Simulations Using the CHARMM36 Additive Force Field. J Chem Theory Comput 12, 405–413, doi:10.1021/acs.jctc.5b00935 (2016).

80 Wu, E. L. et al. CHARMM-GUI Membrane Builder toward realistic biological membrane simulations. J Comput Chem 35, 1997–2004, doi:10.1002/jcc.23702 (2014).

81 Jo, S., Lim, J. B., Klauda, J. B. & Im, W. CHARMM-GUI Membrane Builder for mixed bilayers and its application to yeast membranes. Biophys J 97, 50–58, doi:10.1016/j.bpj.2009.04.013 (2009).

82 Jo, S., Kim, T. & Im, W. Automated builder and database of protein/membrane complexes for molecular dynamics simulations. PLoS One 2, e880, doi:10.1371/journal.pone.0000880 (2007).

83 Lee, J. et al. CHARMM-GUI Membrane Builder for Complex Biological Membrane Simulations with Glycolipids and Lipoglycans. J Chem Theory Comput 15, 775–786, doi:10.1021/acs.jctc.8b01066 (2019).

84 Alvares, D. D. S. et al. Modulatory Effects of Acidic pH and Membrane Potential on the Adsorption of pH-Sensitive Peptides to Anionic Lipid Membrane. Membranes (Basel) 11, doi:10.3390/membranes11050307 (2021).

85 Eastman, P. et al. OpenMM 7: Rapid development of high performance algorithms for molecular dynamics. PLoS Comput Biol 13, e1005659, doi:10.1371/journal.pcbi.1005659 (2017).

86 Huang, J. et al. CHARMM36m: an improved force field for folded and intrinsically disordered proteins. Nat Methods 14, 71–73, doi:10.1038/nmeth.4067 (2017).

87 Humphrey, W., Dalke, A. & Schulten, K. VMD: visual molecular dynamics. J Mol Graph 14, 33–38, 27-38, doi:10.1016/0263-7855(96)00018-5 (1996).

88 Marx, D. C. & Fleming, K. G. Local Bilayer Hydrophobicity Modulates Membrane Protein Stability. J Am Chem Soc 143, 764–772, doi:10.1021/jacs.0c09412 (2021).

89 Fleming, P. J. & Fleming, K. G. HullRad: Fast Calculations of Folded and Disordered Protein and Nucleic Acid Hydrodynamic Properties. Biophys J 114, 856–869, doi:10.1016/j.bpj.2018.01.002 (2018).

90 Fleming, P. J., Correia, J. J. & Fleming, K. G. Revisiting macromolecular hydration with HullRadSAS. Eur Biophys J, doi:10.1007/s00249-022-01627-8 (2023).

91 Heinig, M. & Frishman, D. STRIDE: a web server for secondary structure assignment from known atomic coordinates of proteins. Nucleic Acids Res 32, W500–502, doi:10.1093/nar/gkh429 (2004).

92 Gutzeit, V. A. et al. A fine-tuned azobenzene for enhanced photopharmacology in vivo. Cell Chem Biol 28, 1648–1663 e1616, doi:10.1016/j.chembiol.2021.02.020 (2021).

93 Vivaudou, M. et al. Probing the G-protein regulation of GIRK1 and GIRK4, the two subunits of the KACh channel, using functional homomeric mutants. J Biol Chem 272, 31553–31560, doi:10.1074/jbc.272.50.31553 (1997).

